# A detailed study Biofilm forming Polyextremophilic Firmicutes from the Himalayas and exploration of their plant growth promoting potential

**DOI:** 10.1101/2025.05.29.656832

**Authors:** Bedaprana Roy, Debapriya Maitra, Archisman Chakraborti, Pijush Basak, Indranath Chaudhuri, Jaydip Ghosh, Arup Kumar Mitra

## Abstract

Polyextremophily has been an intriguing point of interest among the scientists, a phenomenon in which organisms can thrive in combinations of multiple extreme conditions. In the foreseeable future, climatic conditions are predicted to change, and agriculture is bound to suffer with these changes in the environment. The situation necessitates some solutions that can ensure agricultural sustainability, in the changed, harsh environmental conditions without more toxic chemical intrusions in the agricultural fields.

The bacterial strains *Brevibacillus parabrevis BRAM_Y3 and Mesobacillus subterraneous BRAM_Y2* in this study, isolated from the waters of Yamunotri showed incredible properties of Polyextremophily like tolerance to not only temperatures as high as 70°C, pH range (4-10), saline, drought, UV and heavy metals such as mercury, iron, silver and arsenic but also their two-dimensional and three-dimensional combinations. The elevated levels of biofilm formation when subjected to stress revealed their property of using biofilm as their shield against harsh conditions. Furthermore, they were studied for their plant growth promotion like nutrient sequestration (N, P, K, plant hormone production etc.) biocontrol properties (cell wall degrading enzymes, siderophores, VOCs etc.) and the ability to confer abiotic stress resistance (ACC Deaminase) to plants. The in-vivo experiments conducted on *Zea mays*. L yielded conclusive and promising results. The use of these polyextremophiles for agriculture in harsh environments may serve as a solution for the global warming mediated climate and environmental changes.

## Introduction

The Garhwal Himalayan glaciers are a hub of microbial biodiversity. Yamunotri [[31.01°N 78.45°E](https://geohack.toolforge.org/geohack.php?pagename=Yamunotri&params=31.01_N_78.45_E_)], is the source of the river Yamuna situated in Champasar glacier, north of Uttarkashi, at an elevation 3293 meters above sea level. The river Yamuna actually originates from a frozen lake, at an elevation of 4421meters, but due to its inaccessibility, the shrine of Yamunotri is located in the foothills of the glacier. The area has a number of hot water springs, the most important of them being the Suryakund from which the water sample for this investigation had been collected. The hot spring water could supposedly have possessed extremophiles, and even polyextremophiles. Polyextremophiles, capable of surviving at multiple forms of extremes might have inhabited the earth during the origin of life and some of them still live on in some specific niches on the surface of the earth. They are also known to serve as analogues of extra-terrestrial life forms that might exist other planets (Seckbach. J, et al., 2013). But, along with all these interesting applications of polyextremophiles, another field they can significantly contribute to is the gradual changing environment due to global warming, that these organisms are already adapted to, the topic addressed in this study. In an investigation by Liverman, 1991, it was explained that future changes in climate will decrease the soil moisture, making it dry which will severely deplete agricultural productivity of the soil. This dryness will further lead to increased requirement of irrigation. The increase in irrigation rates will in turn increase the soil water table leading to significant increase in the soil salinity severely affecting agriculture as well (Utset. A. et. al, 2001). The reduction in soil organic matter is directly related to global warming in a vicious cycle. Organic matter reduction increases the greenhouse gasses, increasing global warming. The temperature increase due to this global warming can then affect the decomposition pattern of the soil organic matter and the cycle goes on (Miko U.F, 1995).

In this alarming situation, polyextremophiles with incredible plant growth promoting and soil enriching qualities can come to the rescue. The microorganisms can thrive in the hostile environment of the soil created by global warming and climate change. Properties like nitrogen fixation will sustain the soil nitrogen content (DeLuca, T.H. et al., 1996) production of ligno-cellulytic enzymes will balance the soil organic carbon (Hemati. A et al., 2021) Production of ACC Deaminase, will help the plants survive the abiotic stresses in the soil (Glick, B. R., 2004). Some special properties such as plant hormone production and siderophore production will also have great impacts on the plant and soil health without inflicting any other harm on the general environment (Kraepiel, A.M.L. et al., 2009).

Reflecting upon this concept, this investigation was designed, where two biofilm-producing bacterial strains were characterised for their polyextremophilic properties which were further associated with their biofilm producing properties. Then these polyextremophilic strains *Brevibacillus parabrevis BRAM_Y3 and Mesobacillus subterraneous BRAM_Y2* were assessed for their plant growth promoting properties, biocontrol properties and properties contributing to environment sustainability. *Brevibacillus parabrevis* has been known to be a gram positive, rod-shaped, obligate aerobic mesophile, with slow growth on broth and heavy growth on agar media. They have been mostly observed to produce endospores. to Its ability in bioremediation has been explored by Ismail S et al., 2023 in response to lead toxicity but it has not been seen as an extremophile previously. Though the *Brevibacillus* genus have been sparsely studied for their plant growth promoting potential, there has been no reports for *Brevibacillus parabrevis* as a plant growth promoting microorganism. But, *Brevibacillus parabrevis* BRAM_Y3 studied in this investigation has been found to be polyextremophilic in nature, with no endospore formation and having enormous activities in context of plant growth promotion.

For, *Mesobacillus subterraneous*, the genus has been identified by Kanso et al., 2002 as gram positive or gram variable, rod shaped bacteria that are either aerobes or facultative anaerobes. But the report does not talk about any extremophilic nature of the bacterial species, and the taxon till then had not been designated as “*Mesobacillus*” which was finally resolved by Patel and Gupta 2020.

These two unique strains in this investigation might play a promising role in agriculture in harsh environmental conditions were using mesophilic plant growth promoting bacterial strains might be rendered useless.

## Materials and Methods

### Sampling and Evaluation of the physical characteristics of the Yamunotri, Suryakund water

The sample of water studied in this investigation was obtained from a hot water spring situated in the Yamunotri Temple complex, Yamunotri, Uttarakhand, India at an altitude of 3,293 metres (10,804 ft) above sea level. The sampling was done in sterile containers and gloves were worn to avoid sample contamination. The container was sealed and transported to the lab for further analysis. The general physical characterization of the water was done using an Elico PE 138 Water Quality Analyzer ( https://www.elico.co/pe-138).

### Isolation, characterization and Identification of Bacterial Strains from Yamunotri, Surya Kund

#### Isolation

The isolation of bacterial strains was carried out via standard serial dilution methods followed by spread plating on Nutrient Agar medium. The two colonies BRAM_Y2 and BRAM_Y3 was isolated on the basis of their prevalence in all the dilution plates.

#### General Characterization of BRAM_Y2 and BRAM_Y3

The pure cultures of the two strains were first analysed for their colony morphologies. Then Gram staining was carried out following established procedures. The endospore staining of the two strains were carried out using the established procedure of the Schaeffer-fulton method. The antibiotic sensitivity test was carried out by the Kirby-Bauer disk diffusion method for both BRAM_Y2 and BRAM_Y3 using *Staphylococcus aureus* as control. The Samples of the *Bacillus* sp. and BRAM Y2 and BRAM Y3 strains were prepared for scanning electron microscopy. Using Omar Farhan Ammar, 2017, as a guide, the laboratory strain was fixed for electron microscopy using the conventional glutaraldehyde procedure. The strains were imaged using a ZEISS EVO 18 variable pressure scanning electron microscope.

#### Molecular identification of BRAM_Y2 and BRAM_Y3

A portion of the 16S rDNA gene was enlarged utilizing the 27F and 1492R primers. Afterward, a distinct PCR amplicon band measuring 1500 base pairs was visualized on an Agarose gel. The PCR product underwent purification to eliminate impurities. Subsequently, both forward and reverse DNA sequencing reactions of the PCR product were executed with the respective primers, utilizing the BDT v3.1 Cycle sequencing kit on the ABI 3730xl Genetic Analyzer. The consensus sequence of the 16S rDNA gene was then derived from the aligned forward and reverse sequences using alignment software. This resultant sequence was subjected to a BLAST search against the NCBI GenBank database, and the top ten sequences with the highest identity scores were selected. These sequences were then aligned using the Clustal omega software to generate a distance matrix. Finally, the MEGA 7 software, was utilized to generate a distance matrix, followed by the generation of a phylogenetic tree, produced with regard to the first 10 sequences. (Felsenstein J.,1985; Kimura M.,1980; Kumar S., 2015)

### Biofilm Formation and subsequent characterization

#### Qualitative Analysis

The qualitative test for analysing biofilm formation by the two strains was performed by streaking the bacterial colonies on Brain heart infusion (BHI) Congo red agar. Biofilm formation on this agar medium is indicated by the development of thick, mucoid and black colonies on the agar plate. (Roy, et.al, 2022)

#### Quantitative determination of formation of biofilm by the strains

The amount of biofilm formation was measured using the tissue culture plate method with 0.1% crystal violet as the stain. The biofilm attached to the plate was then analysed using a Readwell Robonik Elisa reader at 570 nm (Roy, Maitra and Mitra, 2020)

##### Evaluation of the Biochemical Characteristics of the Biofilms produced by the two strains

###### Induction of Primary Biofilm Production

The biofilm produced by BRAM_Y2 and BRAM_Y3 along with *Bacillus* sp. control lab strain was harvested by the help of a similar strategy to Raghad R and Al Abbasi, 2013. The production of the bacterial biofilm was stimulated in basal salt solution, enhanced using 3% glucose maintaining the pH at 7± 0.1. 2ml of 18hour cultures of BRAM_Y2 and BRAM_Y3 strains growing in standard nutrient broth was used to inoculate the aforementioned medium for three days.

###### Isolation of the Biofilm matrix from culture

Centrifugation at 5000 rpm for 10 minutes precipitated the BRAM Y2 and BRAM Y3 and *Bacillus* sp. Control Laboratory strain cells from the 3 days old culture in basal salt solution enhanced with glucose. Then, ethanol (stored at low temperatures) was added in a 1:2 volume ratio. This was followed by the setup being incubated for 24 hours at 4°C to allow the biofilm matrix to precipitate out of the supernatant. The biofilm matrix that had been precipitated out was recovered by centrifugation at a specific rpm of 6000 for 20 minutes at normal temperature, followed by 24-48 hours of full drying at 70°C. Scraped and gathered in a centrifuge tube, the EPS powder. (Raghad R. and Al Abbasi, 2013).

###### Quantitative estimation of the various Biochemical elements of the Biofilm

To determine the specific concentration of, carbohydrates, DNA, and protein in the biofilm formed by each ml of culture, of the biofilms produced by BRAM_Y2 and BRAM_Y3 and *Bacillus* sp. laboratory strain three different protocols were adopted. Utilizing diphenyl amine, Burton’s approach was used to estimate the DNA (Zulfikar Ali et.al, 2014). The conventional Lowry method was used to estimate the protein (Lowry et.al, 1951). Last but not least, (Dubois et al. (1956)) evaluation of the total content of carbohydrate in the biofilm matrix of the two strains using the phenol-sulfuric acid method.

### Verification of extremophilic and Polyextremophilic properties and its correlation with Biofilm formation

#### Single-dimensional stress

##### Temperature tolerance

The growth of BRAM_Y2, BRAM_Y3, and *Bacillus* sp. control Laboratory strain was tested at 3 varied temperatures (70°C, 37°C and 20°C). A small amount (100 micro lit) of a 24-hour old culture of each bacterial strain was placed in a sterile medium in a flask and incubated under shaking conditions. The growth was monitored at regular intervals for 8 hours using a colorimeter at a specific wavelength (600nm), and the results were plotted on a growth curve for each temperature. This method has been described in Roy et.al, 2022.

##### Thermal death time and Cold shock treatment

The thermal death time of the bacterial cultures was determined by incubating them at a high temperature (120°C) and measuring the number of viable cells over time. For the cold shock treatment, the bacterial cultures were incubated at a low temperature (−20°C) overnight and the growth was monitored at regular intervals to observe any reduction.

##### Salt tolerance

Sodium chloride was added to the bacterial culture medium in concentrations ranging from 1-8%, with 0.5% NaCl serving as the control setup. Along with the bacterial strains BRAM_Y2 and BRAM_Y3, *Bacillus* sp., control laboratory strain cultures were then used. The prepared medium was inoculated with the cultures and then incubated for 24 hours at 37°C. After incubation, optical densities at 600 nm were recorded. (Sarita Kumari., *et.al*, 2019).

##### pH tolerance

Bacillus sp. laboratory strain, BRAM Y2 and BRAM Y3 bacterial cultures grown in different pH in a buffered liquid nutrient medium to maintain the pH. The growth was measured after 24hours spectrophotometrically at 600nm. (Roy et al., 2022)

##### Drought tolerance

To mimic the conditions of drought, the growth medium was amended with Polyethylene glycol 6000 at different concentrations (0%–20%). Along with BRAM_Y2 and BRAM_Y3, *Bacillus* sp., control laboratory strain cultures were inoculated in the enhanced medium. After incubating the laboratory strain in the altered medium for five days at 28 +°C in a shaker, the optical densities at 600 nm were determined. (Sarita Kumari *et.al*, 2019)

##### UV tolerance

BRAM_Y2, BRAM_Y3, and Bacillus sp. control Laboratory strain growth, in presence of ultraviolet rays was measured by inoculating them into fresh nutrient broth and exposing the tubes to UV light (Short UV of 254nm wavelength) of a specific intensity of 11µW/cm^2^ for a set amount of time. The optical densities at a specific wavelength were recorded at regular intervals for a set amount of time (1hour), and the results were plotted on a growth curve showing the optical density over time. (Cayron and Lesterlin, 2019)

##### Heavy Metal Tolerance

For iron, mercury, silver, and arsenic, heavy metal tolerance setups were created. To adjust the bacterial growth medium and create the necessary amounts of the individual metals, the corresponding metal salts were added. The medium was supplemented with ferrous sulphate (FeSO4) to supply iron. Silver nitrate (AgNO3), mercuric chloride (HgCl2), sodium asenite (NaAsO2), and silver (Ag) were used to assess the tolerance to mercury (As). A sterile broth with the required metal amendments was kept for each metal concentration to serve as a control. The Bacillus sp. Laboratory strain and the bacterial strains BRAM Y2 and BRAM Y3 were added to the media that had been altered with the four metal salts stated above in accordance with different metal concentrations. The media were incubated at 37°C with shaking for one night. The bacterial growth, as indicated by the optical density (OD), was then measured at 600 nm using the corresponding sterile medium as a reference.” (Roy *et.al,* 2022).

#### Two-dimensional stress

The growing ability of BRAM_Y2, BRAM_Y3, and *Bacillus* sp. control Laboratory strain was tested under 16 different combinations of 2D stress conditions. The combinations were (Temperature 70 °C + 20% PEG, Temperature 70 °C + 4% NaCl, Temperature 70 °C + As 100, Temperature 70 °C + pH4, Temperature 70 °C + pH10, pH4+4% NaCl, pH10 + 4% NaCl, UV+20% PEG, UV + As 100 ppm, UV + Fe 200 ppm, UV + pH4, UV + pH10, Temperature 20 °C + Fe 200 ppm, Temperature 20 °C + 4% NaCl, Temperature 20 °C + pH4, and Temperature 20 °C + pH10). The biofilm formation of these stressed samples was then measured using the standard tissue culture plate method and 0.1% CV stain. The results were plotted and compared to the control, which was a no-stress condition. (Roy et.al, 2022)

#### Three-dimensional stress

The growing ability of BRAM_Y2, BRAM_Y3, and *Bacillus* sp. control Laboratory strain was tested under 9 different combinations of 3D stress conditions in broth cultures. The combinations were (Temperature 20 °C + 20% PEG+4% NaCl, Temperature 20 °C + pH10 + 4% NaCl, Temperature 20 °C + pH10+Fe200 ppm, pH10 + 4% NaCl+20% PEG, Temperature 70 °C + pH10+As100 ppm, Temperature 70 °C + pH10 + 4%NaCl, Temperature 70 °C + 4% NaCl+20% PEG, UV+4% NaCl+20% PEG and UV + Temperature 20 °C + 20% PEG). After being incubated overnight, the biofilm formation of these stressed samples was measured using the crystal violet staining method on tissue culture plates, with 0.1% CV as the stain. The results were plotted and compared to the control, which was a no-stress condition. (Roy et.al, 2022)

### Study of the changes in dynamics of bacterial biofilm with 3d stress using Fourier Transform Infrared Spectroscopy

The bacterial biofilm was isolated from a 3D stress setup (combination (Temperature70 °C + 4% NaCl+20% PEG) that showed best growth) and no stress setups using the methodology described in earlier. The study was carried out using KBr as the carrier in FTIR (model: Bruker Alpha II; Made in Germany).

#### Application in Agriculture and Environment sustainability

##### Plant Growth Promotion Abilities

###### Nitrogen fixation

The nitrogen fixation ability of the two strains BRAM_Y2 and BRAM_Y3 along with *Bacillus* sp. control Laboratory strain was checked by plating them on modified Jensen’s Agar media with indicator Bromothymol blue. The change in the colour of the medium after incubation indicated the different stages of nitrogen fixation. (Sulistiyani T, & Meliah S., 2017)

###### Phosphate solubilization

For the determination of Phosphate solubilising ability of BRAM_Y2 and BRAM_Y3 along with *Bacillus* sp. control Laboratory strain, the strains were plated in Pikovskaya Agar medium (Hi-media) amended with the indicator Bromothymol blue. (Maitra et.al, 2022)

###### Potassium solubilization

Potassium solubilization was detected my plating the two strains BRAM_Y2 and BRAM_Y3 along with *Bacillus* sp. control Laboratory strain On Aleksandro Agar medium modified with acid base indicator Bromothymol blue. The use of the indicator clearly depicted the production of different acids during the process of solubilization. (Maitra et.al, 2022)

###### Indole Acetic Acid

The IAA production by BRAM_Y2 and BRAM_Y3 along with Bacillus sp. Laboratory strain was estimated using the Salkowski reagent as per Sarker and Rashid, 2013. The measurement was done in two set ups, first with the IAA precursor, 0.1% Tryptophan supplemented in the culture medium and the second without the precursor.

###### Gibberellins

Gibberellic acid production by BRAM_Y2 and BRAM_Y3 along with *Bacillus* sp. Laboratory strain was done by using modified Holbrook method by Sharma and Sharma et.al, 2018. (Hollbrook et.al, 1961)

###### Stress Response: ACC Deaminase

Qualitative evaluation of ACC Deaminase: 1-aminocyclopropane-1-carboxylic acid deaminase production was checked by plating BRAM_Y2 and BRAM_Y3 along with *Bacillus* sp. Laboratory strain in Nitrogen free Dworkin foster medium, with 3mM ACC supplement (Sigma Aldrich) as the sole nitrogen source. Ammonium chloride was used in one setup as the positive control and minimal Dworkin foster medium devoid of nitrogen source was used as the negative control. (Kumar et.al, 2012)

Quantitative Estimation of ACC Deaminase: Quantification of ACC Deaminase production by BRAM_Y2 and BRAM_Y3 along with *Bacillus* sp. Laboratory strain was performed using the ninhydrin assay method as per Li Z. et al., 2011.

###### Qualitative estimation of Siderophore production

The CAS agar diffusion assay or CASAD for BRAM_Y2 and BRAM_Y3 along with *Bacillus* sp. Laboratory strain was performed using a modified Schwyn and Neilands, 1987. The modified method was adapted from Shin et al., 2000.

###### Quantitative evaluation of Siderophore production

Quantification of Siderophore production by BRAM_Y2 and BRAM_Y3 along with *Bacillus* sp. Laboratory strain was also done by the traditional method from Arora and Verma, 2017. The measurement of siderophore production was done in percent siderophore unit (psu) with the following formula:

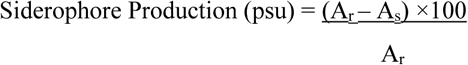

Where, A_r_ is the optical density of CAS solution and un-inoculated broth at 630nm and A_s_ is the optical density of the sample, CAS solution and cell free supernatant of the bacterial; culture. (Payne, 1993)

###### Typing of the Siderophore produced by BRAM_Y2 and BRAM_Y3

The bacterial strains BRAM_Y2 and BRAM_Y3 along with *Bacillus* sp. Laboratory strain were checked for two types of siderophore, hydroxamate and catecholate. The hydroxamate type was detected by the tetrazolium test and the catecholate type was detected by the Arnow’s test following Radhakrishnan et al.,2014.

##### Biocontrol Properties of BRAM_Y2 and BRAM_Y3

###### Peroxidase(total)

To measure the total peroxidase production of BRAM_Y2, BRAM_Y3, and *Bacillus* sp. control Laboratory strain, the cultures were centrifuged and the cell pellets were suspended in a buffer (20 mM sodium phosphate buffer; pH 7.4). The cell suspension was then sonicated for a set amount of time (3 seconds) using a specific output (Amplitude of 50) and strokes (7), and the resulting homogenate was centrifuged under cold conditions. The supernatant was then used as the crude enzyme extract for the peroxidase assay, following the method described in Kalyani et al. (2011).

###### Catalase

The catalase production by BRAM_Y2 and BRAM_Y3 along with *Bacillus* sp. control Laboratory strain was quantified using 30% Hydrogen peroxide solution following the method of Iwase T. et al., 2013.

###### Protease

For the quantification of Protease production by BRAM_Y2 and BRAM_Y3 along with *Bacillus* sp. control Laboratory strain, the bacterial strains were inoculated in MRS broth for 10days and centrifuged to get the crude enzyme extract. Enzyme quantification was carried out by the method of Tsuchida et al. (1986) using Casein as a substrate.

###### Pectinase

Cell free supernatant of BRAM_Y2 and BRAM_Y3 along with *Bacillus* sp. Laboratory strain was utilised as the crude enzyme extract for quantification of pectinase production by the strains. The assay was carried out using 1% (w/v) Pectin as substrate following the methodology by Shrestha et al., 2021.

###### Beta-1,3-glucanase

To quantify the production of Beta-1,3-glucanase by BRAM_Y2 and BRAM_Y3 along with *Bacillus* sp. control Laboratory strain, 1% laminarin was used as the substrate followed by the measurement of the reducing sugars formed by the enzymatic reaction by DNS method. (Rais A. et al., 2017).

###### HCN

HCN Production by BRAM_Y2 at7 BRAM_Y3 along with Bacillus sp. Laboratory strain was checked by streaking the bacterial 5olonies on Kings B agar medium with 4.4g/l glycine amendment. A filter paper dipped in Picric acid solution was placed in the upper lid of the petri dish. The dishes were sealed with parafilm and incubated at 28°C for 48 hours and the change in colouration of the filter paper was observed. (Reetha A.K et al., 2014)

###### Ammonia

The Ammonia production by BRAM_Y2 and BRAM_Y3 along with Bacillus sp. Laboratory strain was detected by first inoculating the bacteria in peptone broth, incubated for 7 days at 30°C followed by the addition of Nesseler’s reagent. The brown colour development indicated production of ammonia.

###### Interaction with Phytopathogenic fungi

The bacterial strains BRAM_Y2 and BRAM_Y3 were spread plated in a nutrient agar plate, and a phytopathogenic fungus, *Fusarium fujikuroi (GenBank Accession: OR426451.1*) isolated from *Zea mays*, whose mycelial disk was placed in a potato dextrose agar plate. The lids of both these plates were discarded and the bottoms were sealed with parafilm facing each other so as to create a physical barrier between the fungus and the bacteria, yet maintain them in the same micro-environment. Thus, ensuring that, any sort of inhibition of fungal growth that takes place after this will be due to the action of the VOCs secreted by the bacteria under experimentation. ((Ruangwong et al., 2021)

##### Field Application on *Zea mays* L

###### Treatment application and Data collection

The bacterial strains BRAM_Y2 and BRAM_Y3 were applied individually and together in a consortium, on test crop *Zea mays* (Variety: KOHINOOR 595). Three plants were taken per setup. The experiments were conducted in the month of September to November for sole crop which is a favourable time for maize harvest as per ICAR guidelines. Cell pellets dissolved in distilled water was used as the mode of treatment. 7.5ml of 24 hours old bacterial suspension (the optical densities of the cultures were 0.7-0.8 at OD_600nm_ containing 9.8 * 10^9 to 11.2 * 10^9 CFU per ml of bacterial culture), diluted to 15ml with distilled was applied to each plant in the rhizosphere region in a circular pattern. The first treatment was applied at the end of 3^rd^ week and then 2 more treatments were applied at a 2-week interval, for a total number of three times in the 90 days lifespan of the plant (Roy et.al., 2023). Regular watering was carried out as per the standard parameters of *Zea mays* cultivation by International Maize and Wheat improvement centre (CIMMYT). Physical data was collected at 1week interval with respect to a control where no treatment was applied. The yield parameters were recorded once the fruiting was complete, such as the “total number of fruits”, “cob weight”, “cob length”, “100 seed weight” and “percent grain filling”.

###### Plant Pigments

Plant pigments such as chlorophyll and carotenoids were measured from the plant leaves after the completion of the treatment application. Acetone extract of leaves was prepared by crushing 0.1gm leaves in 10ml of 80% acetone, followed by centrifugation. The optical densities were measured at 480nm, 645nm and 663nm. This was followed by the calculation of Chlorophyll A, Chlorophyll B and Total Chlorophyll and Carotenoids using the following formula (Kumari. R et al., 2018):

Total Chlorophyll (mg/ g tissue): 20.2(A645) + 8.02(A663)
Chlorophyll a (mg/ g tissue): 12.7(A663) – 2.69(A645)
Chlorophyll b (mg/ g tissue): 22.9(A645) – 4.68(A663)
Carotenoid (mg/ g tissue): [A480 +(0.114(A663) - (0.638-A645)] ×V/1000×W

###### Statistical analysis of the subsequent data

Analysis of variance was measured for the parameters that were taken into account using python 3.11.

##### Environment sustainability

###### Residual Toxicity Enzyme: Urease

The bacterial cultures BRAM_Y2 and BRAM_Y3 along with *Bacillus* sp. Laboratory strain was inoculated in Stuart’s broth. 96-well microtitre plate was used for the assay. The absorbances were measured at 430 nm and 560 nm. The hydrolysis of urea resulted in pink coloration in course of time. The quantification was performed by observing the rate of change of color.

Rate of colour change was measured by the given formula (Okyay and Debora, 2013):

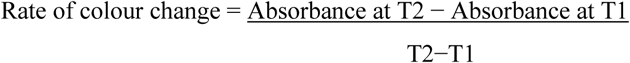

###### Cellulase

Quantification of cellulase production BRAM_Y2 and BRAM_Y3 along with *Bacillus* sp. Laboratory strain was carried out using 0.5% Carboxymethyl cellulose as substrate and cell free supernatant as the crude enzyme extract. The formation of reducing sugars was then measure by using DNS reagent following Sethi S. et al., 2013.

###### Laccase

The production of laccase by BRAM_Y2, BRAM_Y3, and *Bacillus* sp. control Laboratory strain was quantitatively measured using a specific method described in Fatemeh Sheikhi et al., 2012.The enzyme activity was determined by measuring the absorbance at a 465 nm wavelength using a specific substrate (2 mM Guaiacol) in a phosphate buffer at a certain pH (50mM, pH: 6.5). The units of enzyme activity were then calculated and expressed as units per litre, based on the 0.001 unit/min enzyme required to increase the absorbance over time at a certain temperature (55 °C.).

###### Amylase

The production of amylase by BRAM_Y2, BRAM_Y3, and *Bacillus* sp. control Laboratory strain was measured using the cell-free supernatant from the centrifuged bacterial cultures as the crude enzyme extract and starch as the substrate. The enzyme activity was assessed by determining the amount of reducing sugar produced through the breakdown of the substrate using the DNS method. (Patel N. et al., 2019).

###### Improvement in the Soil Health Parameters after application

The physical properties of the soil sample were measured before and after cultivation using a specific method, and the pH and electrical conductivity were measured using corresponding meters. The total organic carbon of the soil was measured using a standard method of wet oxidation by Walkley and Black, and the soil samples were also tested for the availability of various macro and micro nutrients using a specific method. (Piper, C.S. 1966; Jackson,M.L, 1967; Tandon, 1993)

## Results

The water sample in this investigation collected from hot water spring situated in the Yamunotri Temple complex, Yamunotri, Uttarakhand, India, had a slightly alkaline pH varying from 7.9 to 8.1. The TDS was found 6.4 mg/l and the dissolved oxygen was found to be 1.5mg/l. The two isolated strains BRAM_Y2 and BRAM_Y3 grew on nutrient agar medium with specific colony characteristics. BRAM_Y2 had yellowish orange translucent colonies, that were round and elevated (Fig I(a)). BRAM_Y3 on the other hand had opaque white colonies, elevated and round, having a characteristic pearl-like shine (Fig 1(a)). BRAM_Y2 strain was found to be gram variable and BRAM_Y3 was found to be gram positive on gram staining and did not show formation of endospore, when stained for the same. Both the bacterial strains were found to be sensitive to common antibiotics (Table I). The Scanning electron micrograph showed long rod-shaped bacterial structure for both the bacterial isolates which indicated that they could have been *Bacillus* sp. The 16srRNA sequencing identified the BRAM_Y2 as *Mesobacillus subterraneous* BRAM Y2 (Accession number: MW002419) and BRAM_Y3 as *Brevibacillus parabrevis* BRAM_Y3 (Accession number: MW081864) (Fig II).

**Fig I:**
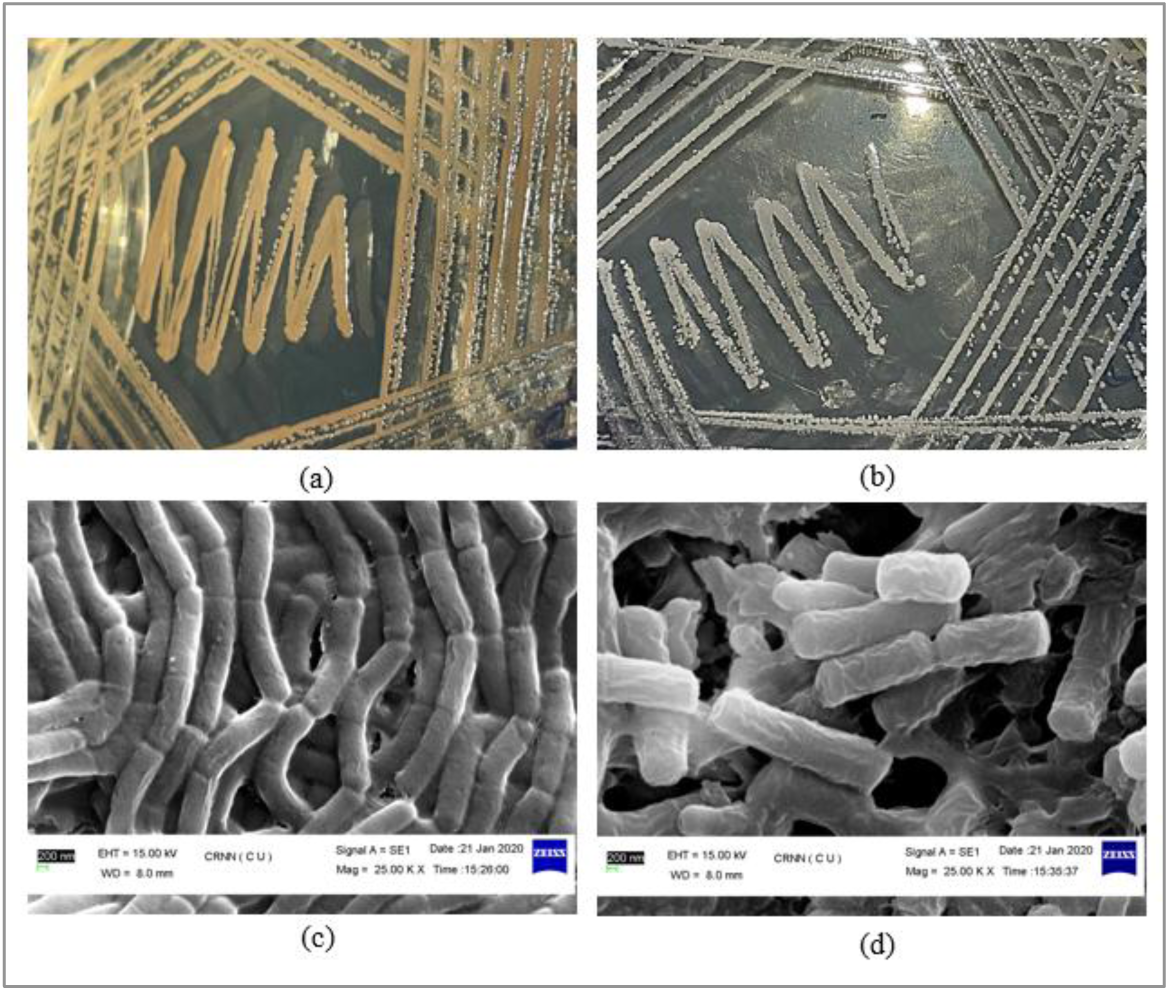
(a) Depicts the colony morphology of BRAM_Y2, with elevated, round and light orange pigmented colonies. (b) Depicts the colony morphology of BRAM_Y3 with round elevated white colonies with characteristic pearl like shine. (c) and (d) Depicts the Scanning electron microscopic image of BRAM_Y2 and BRAM_Y3 respectively, showing rod shaped cells associated with slimy biofilm.

**Fig II.**
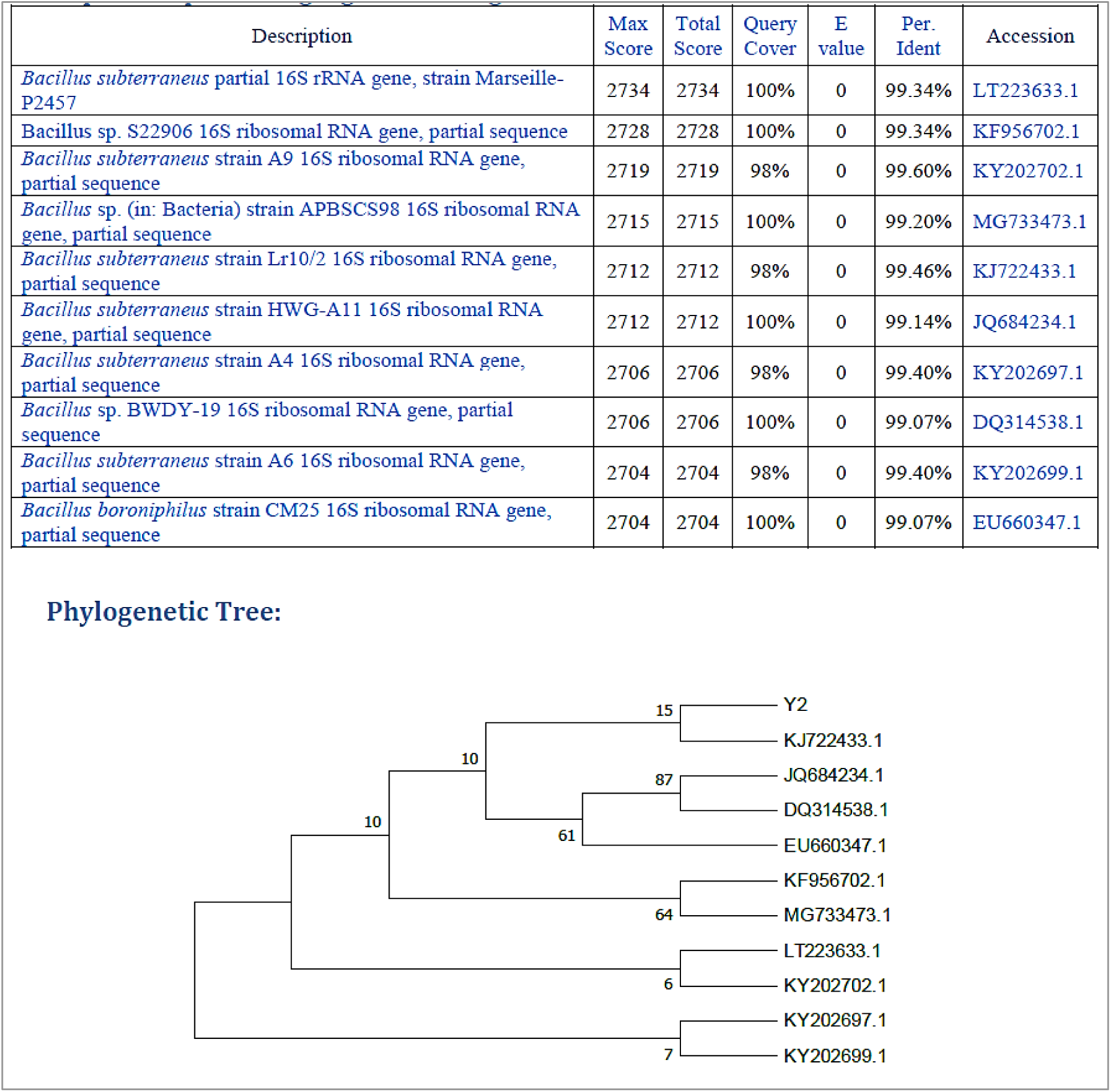
**(a):** The top 10 hits from NCBI Blast results and the phylogenetic tree for BRAM_Y2 16srRNA consensus sequence constructed to understand their evolutionary relationships with the same.

**Fig II.**
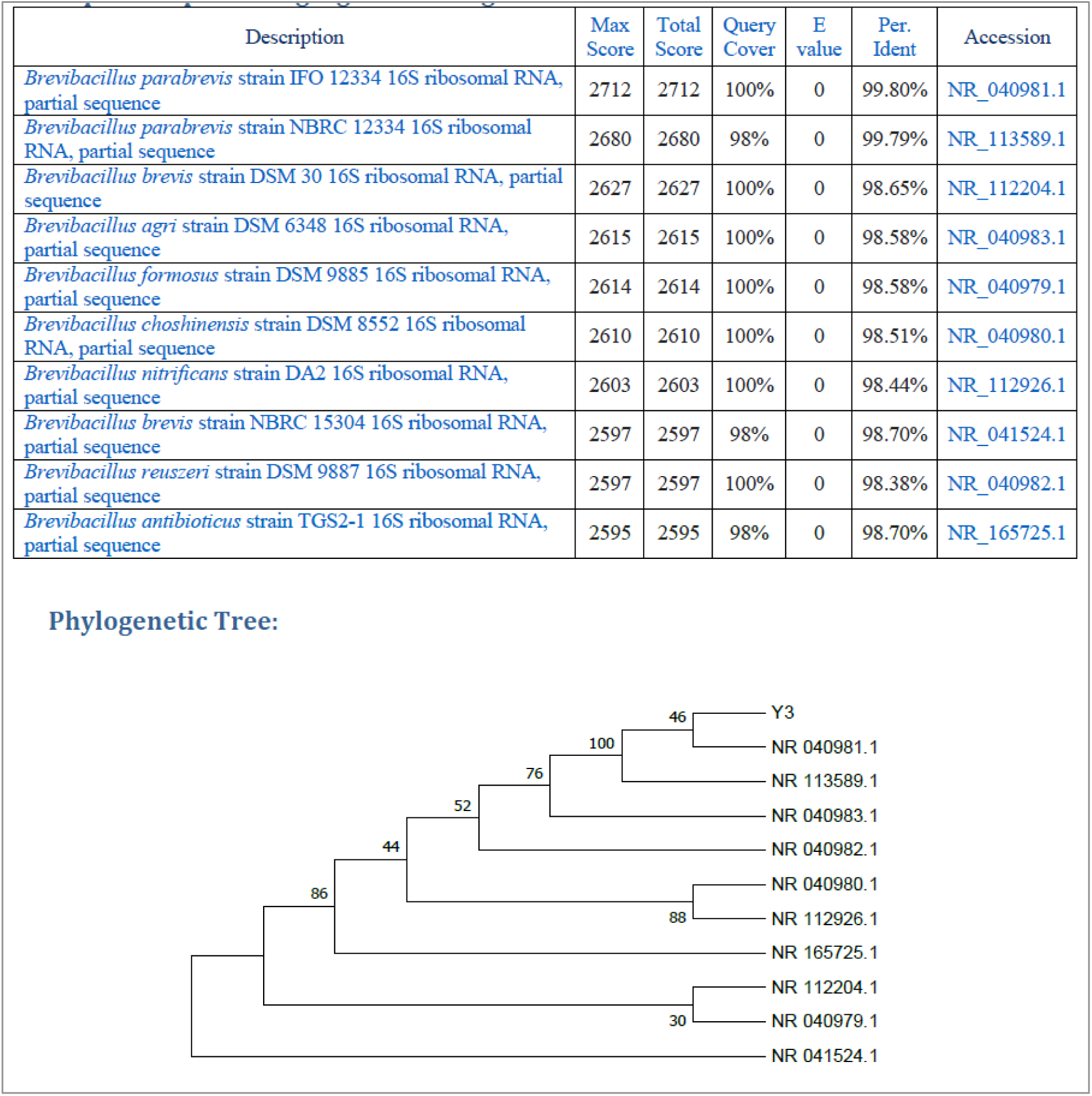
**(b):** The top 10 hits from NCBI Blast results and the phylogenetic tree for BRAM_Y3 16srRNA consensus sequence constructed to understand their evolutionary relationships with the same

**Table I:**
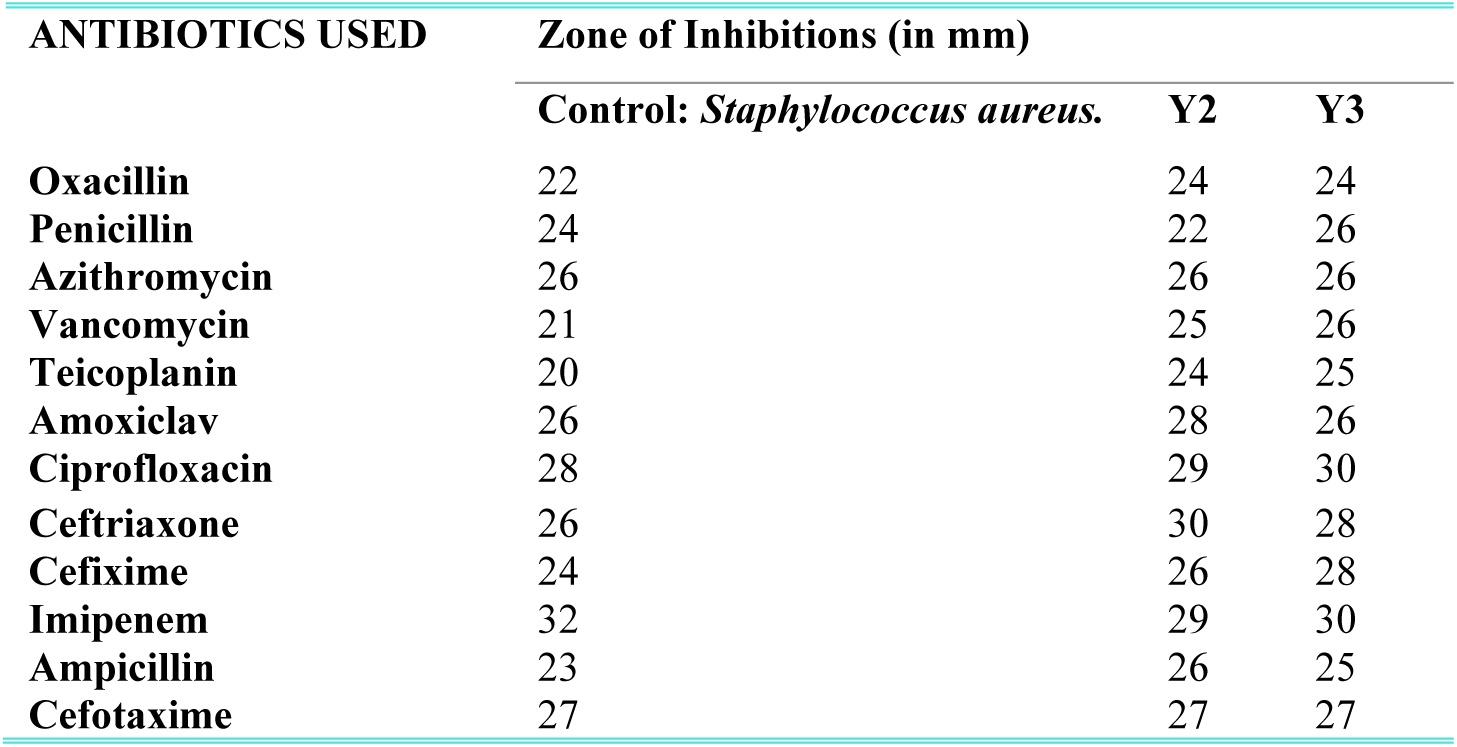
The table shows the antibiotic sensitivity test results of BRAM_Y2 and BRAM_Y3, with the zones of inhibition in mm.

| ANTIBIOTICS USED | Zone of Inhibitions (in mm) |  |  |
| --- | --- | --- | --- |
|  | Control: *Staphylococcus aureus*. | Y2 | Y3 |
| Oxacillin | 22 | 24 | 24 |
| Penicillin | 24 | 22 | 26 |
| Azithromycin | 26 | 26 | 26 |
| Vancomycin | 21 | 25 | 26 |
| Teicoplanin | 20 | 24 | 25 |
| Amoxiclav | 26 | 28 | 26 |
| Ciprofloxacin | 28 | 29 | 30 |
| Ceftriaxone | 26 | 30 | 28 |
| Cefixime | 24 | 26 | 28 |
| Imipenem | 32 | 29 | 30 |
| Ampicillin | 23 | 26 | 25 |
| Cefotaxime | 27 | 27 | 27 |

When these two strains where plated on BHI congo red agar, they showed mucoid black colonies that indicated biofilm formation. The quantification data showed around 15 times greater biofilm formation that the control laboratory strain (Fig III(a)) and as the strains were of rare *Bacilli*, further analysis of the composition of their biofilms were also carried out. The DNA, protein and carbohydrate content of the biofilms of BRAM_Y2 and BRAM_Y3 were analysed and quantified (Fig III(b)).

**Fig III.**
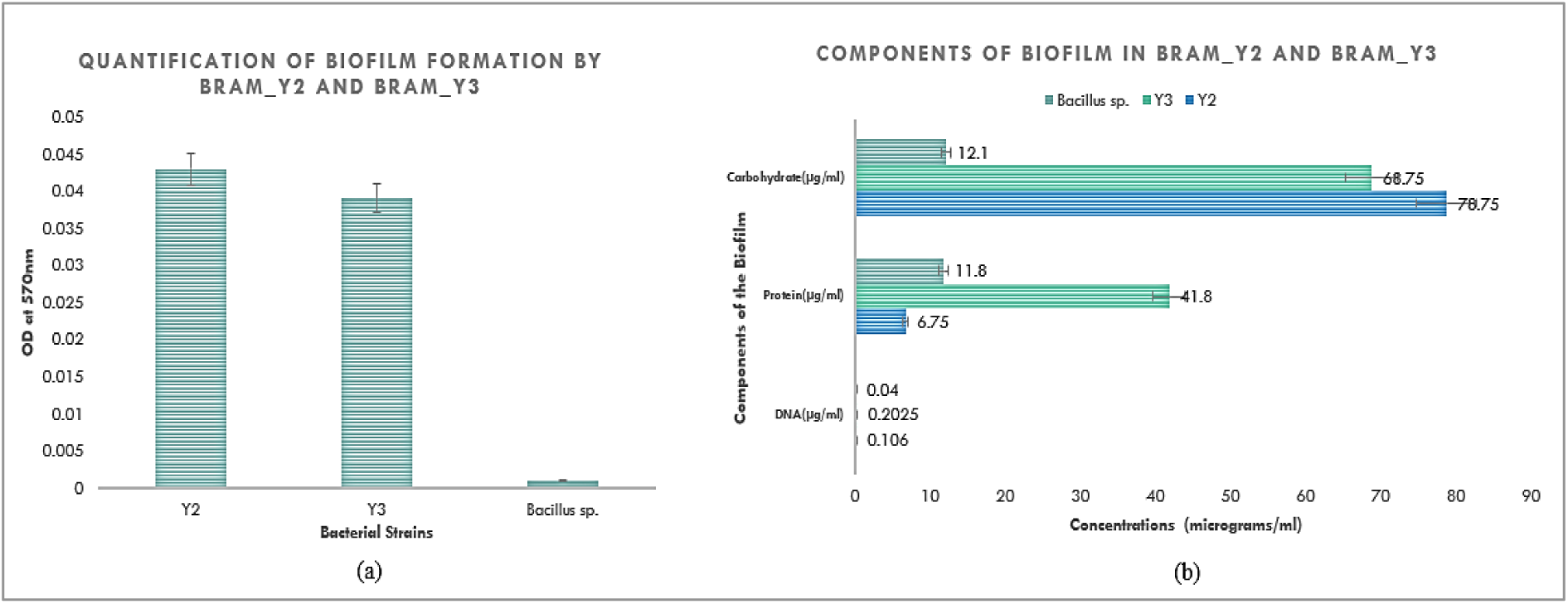
(a) Graphical representation of the quantified results of biofilm formation by BRAM_Y2 and BRAM_Y3. (b) Graphical representation of the quantitative estimation of the biochemical components (DNA, Protein and Carbohydrate) of BRAM_Y2 and BRAM_Y3 biofilm.

Both the bacterial; strains were subjected to 3 temperatures, 20°C, 37°C and 70°C and their growth curves showed their ease of growth in all the three temperatures (Fig IV-(a) and IV(b)). The thermal death was studied at a temperature of time 120 °C and the reduction in the viable cell count was found to be 89%. In the cold shock treatment study, an exposure to −20 °C for 48 hours had the viable cell counts reduced by 74%. To understand the extremophillic nature of the two bacterial strains BRAM_Y2 and BRAM_Y3, they were subjected to different single dimensional stresses and their growth patterns were observed in those extreme environments. The strains showed tolerance and growth up to 4% of NaCl, pH tolerance was found to be from 1 to 12, though proper growth was found in the range of 3-10, with maximum at 10. Drought tolerance was found to be till 20% of Polyethylene glycol concentrations. The bacterial grew luxuriantly at a high dose of UV of 396J/m^2^ in the very first hour. (Fig V). The bacterial strains were also found tolerance to high heavy metal concentrations, of 200ppm Iron, 300ppm pf Arsenic, 20ppm pf silver and 10ppm and 15ppm pf mercury (Fig VI). To conclusively prove the polyextremophilic nature of the two bacterial strains, along with single dimensional stresses they were also subjected to 16 combinations of two-dimensional stresses and 9 combinations of three-dimensional stresses. In addition to that, as the two bacterial strains were exceptional biofilm producers, it was hypothesised that the biofilm might have a significant role in protecting the bacterial strains from the stressed environmental conditions, and thus the increase in biofilm formation when subjected to the 2D and 3d stresses were also recorded (Fig VII, VIII and IX). Plate count was done after each experiment concerning stress so as to keep a record of the bacterial viable cell count which was maintained at 10^7 to 10^8 CFU per ml.------

**Fig IV.**
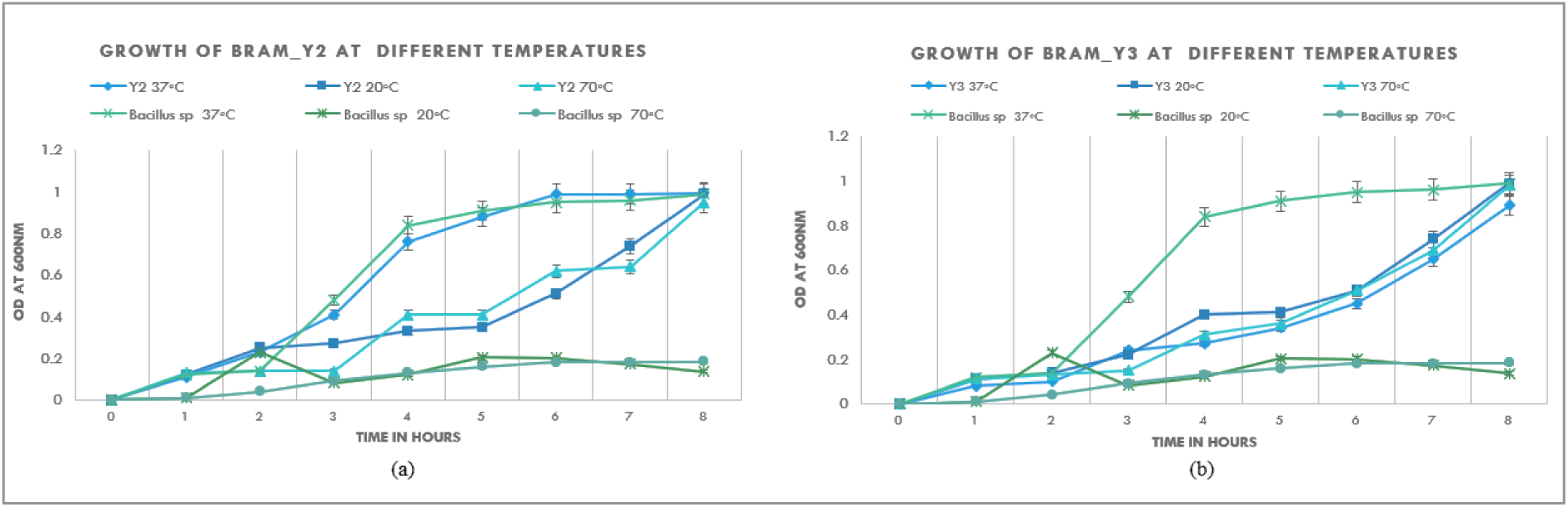
(a) and (b) represents the growth pattern of BRAM_Y2 and BRAM_Y3 respectively at temperature 20°C, 37°C and 70°C with respect to a laboratory strain *Bacillus* sp. as control.

**Fig V.**
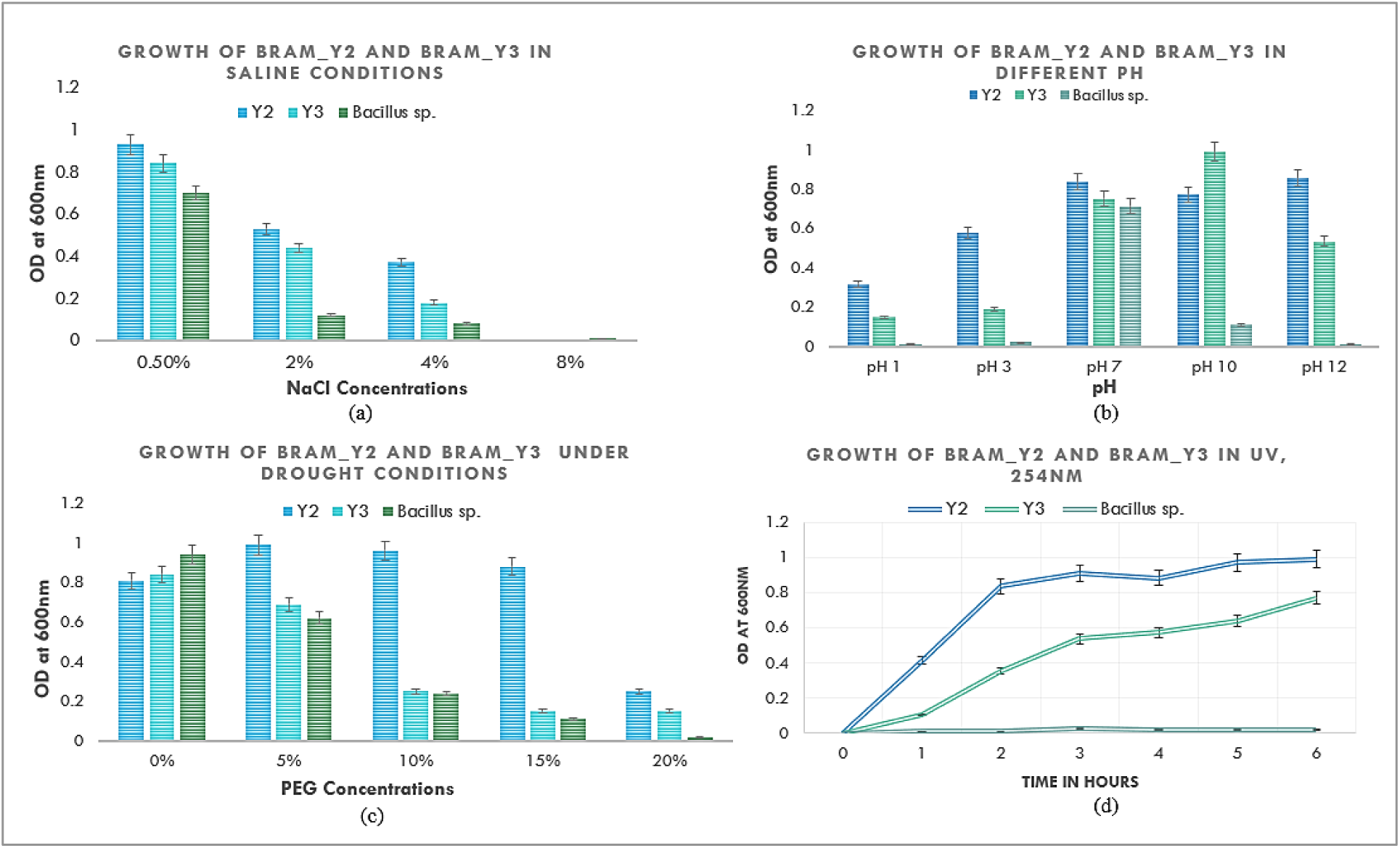
(a) Graphical representation of the growth of BRAM_Y2 and BRAM_Y3 in increasing concentrations of salt (NaCl). (b) Graphical representation of the growth of BRAM_Y2 and BRAM_Y3 in increasing pH from 1 to 12. (c) Graphical representation of the growth of BRAM_Y2 and BRAM_Y3 in increasing concentrations of Polyethylene Glycol 6000 (PEG). (d) Graphical representation of the growth pattern of BRAM_Y2 and BRAM_Y3 in UV light with intensity 11µW/cm^2^, 254nm wavelength.

**Fig VI.**
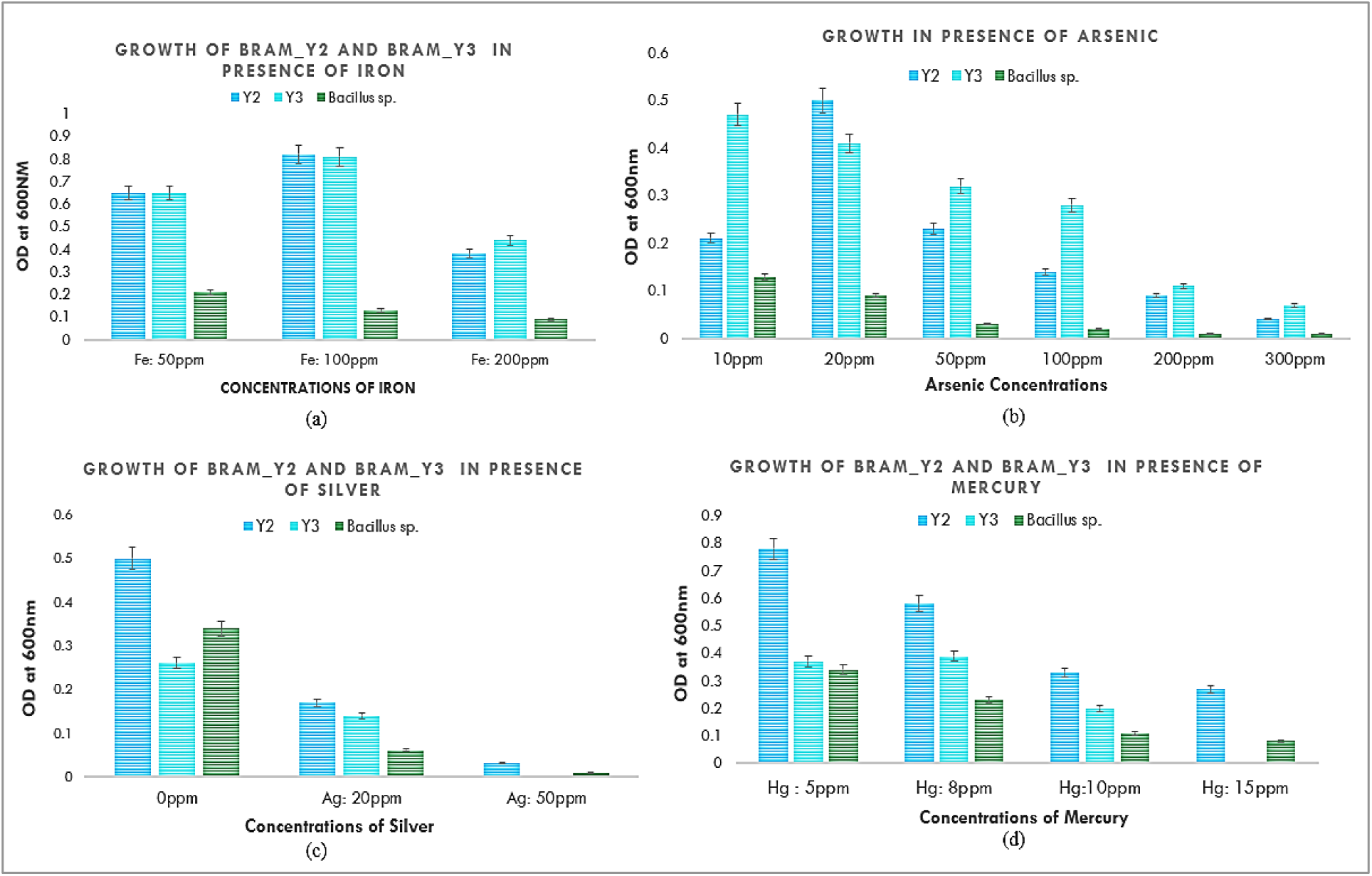
(a) Graphical representation of the growth of BRAM_Y2 and BRAM_Y3 in increasing concentrations of Iron (20ppm to 200ppm). (b) Graphical representation of the growth of BRAM_Y2 and BRAM_Y3 in increasing concentrations of Arsenic (10ppm to 300ppm) (c) Graphical representation of the growth of BRAM_Y2 and BRAM_Y3 in increasing concentrations of Silver (20ppm to 50ppm). (d) Graphical representation of the growth of BRAM_Y2 and BRAM_Y3 in increasing concentrations of Mercury (5ppm to 15ppm).

**Fig VII.**
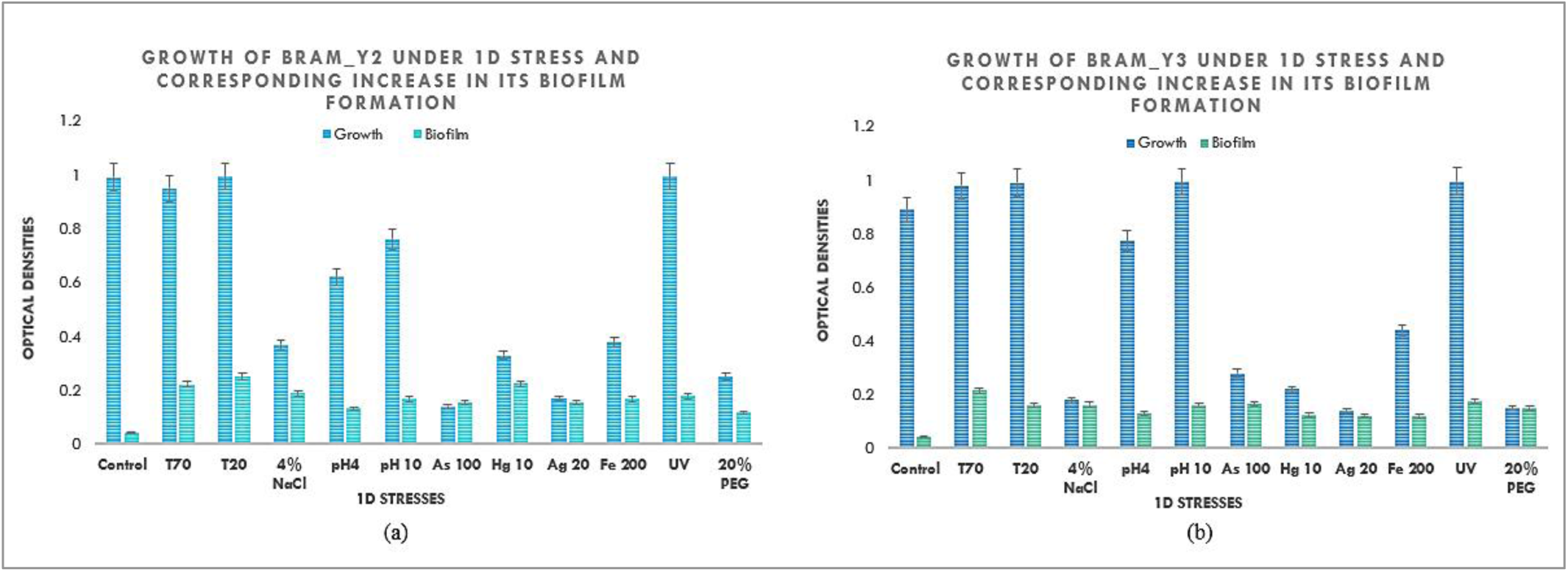
(a) and (b) are the graphical representation of the growth of BRAM_Y2 and BRAM_Y3 respectively in Single dimensional stresses with no-stress control and their corresponding increase in Biofilm formation.

**Fig VIII.**
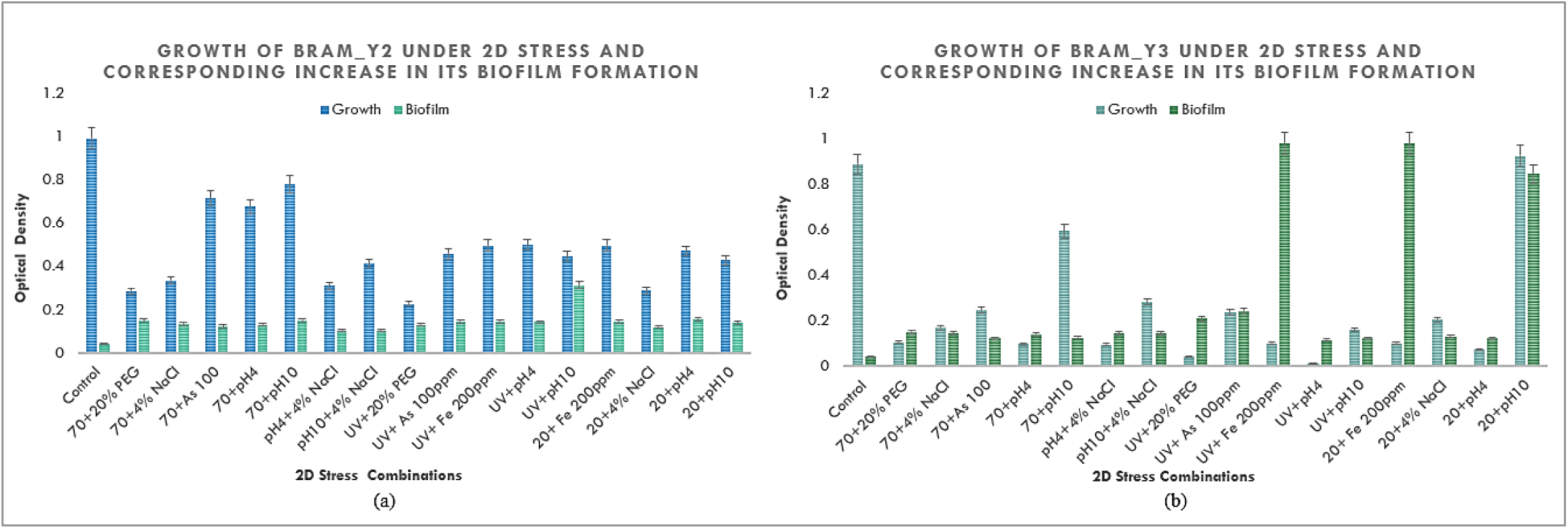
(a) and (b) are the graphical representation of the growth of BRAM_Y2 and BRAM_Y3 respectively in Two-dimensional of stresses with no-stress control and their corresponding increase in Biofilm formation.

**Fig IX.**
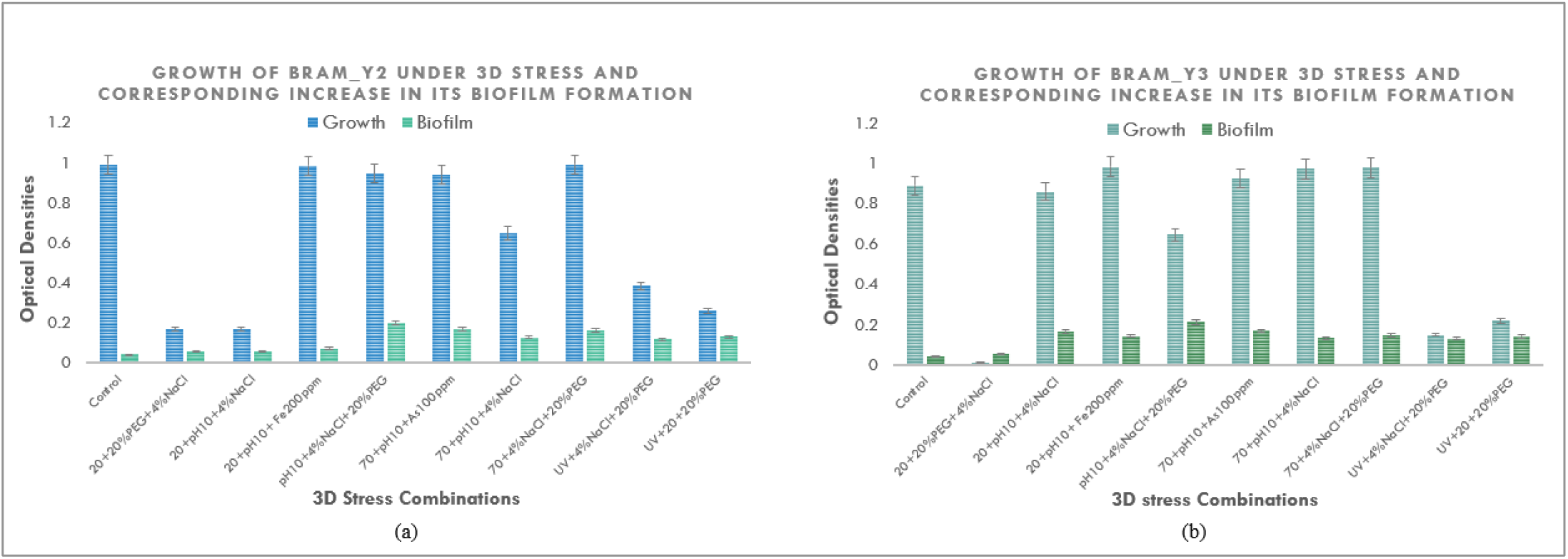
(a) and (b) are the graphical representation of the growth of BRAM_Y2 and BRAM_Y3 respectively in Three-dimensional of stresses with no-stress control and their corresponding increase in Biofilm formation.

After observing increased biofilm formation after stress application, the biofilm was again isolated from “no stress” control and 3D stress combination (Temperature70 °C + 4% NaCl+20% PEG) that showed best growth, and FTIR study was carried out. The spectra showed presence of lipids, proteins, amides I and II (proteins) and some metal-oxygen bonds, but the M-O peaks differed in the 2 strains. The main difference that was observed was in the abundance of the compounds (lipids, proteins etc.). The application of stress increased the abundance of lipids, proteins, amides I and II and tetrahedral metal-oxygen compounds in the 2 bacterial strains. (Fig X).

**Figure X.**
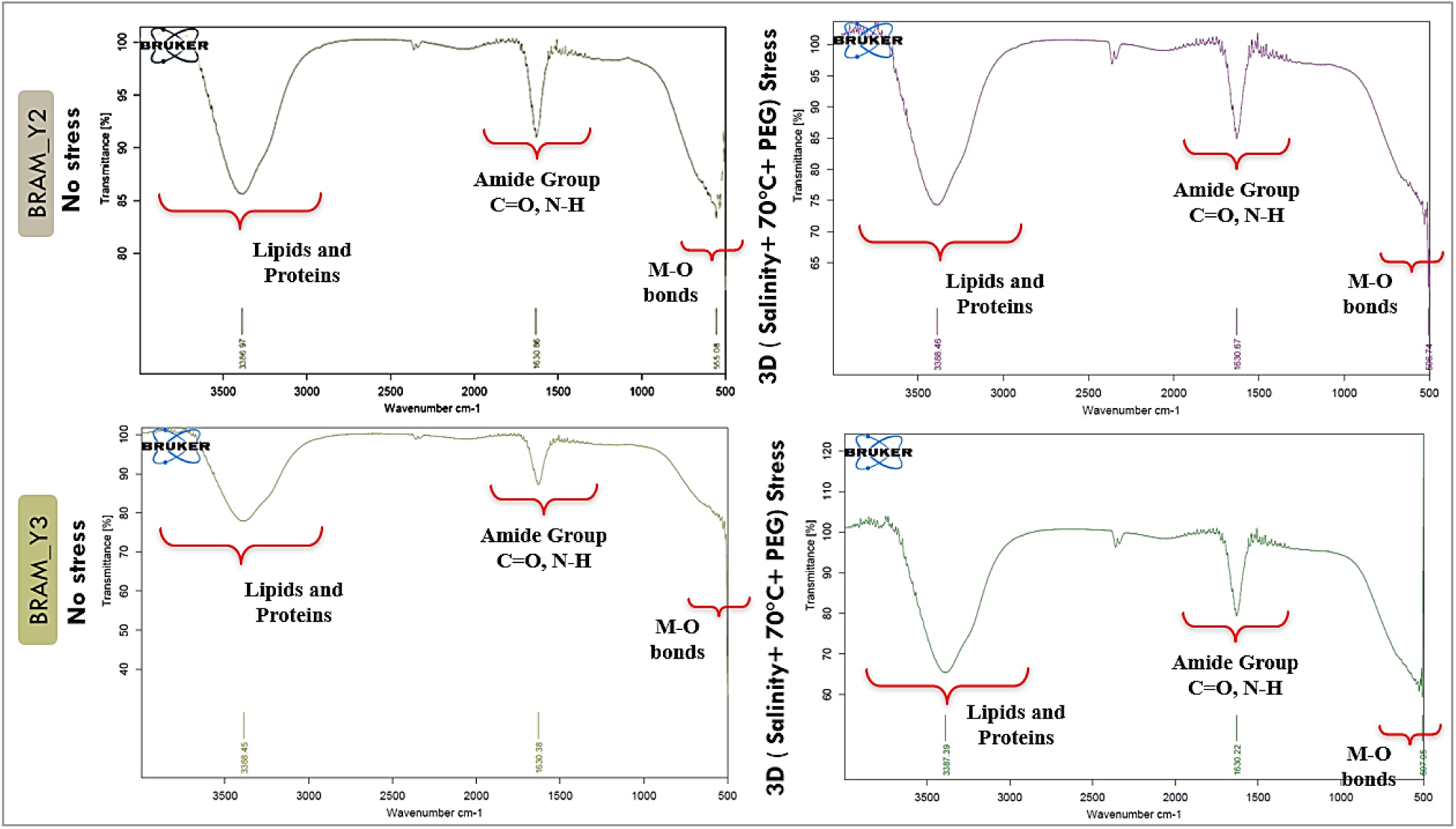
: FTIR studies of the BRAM_Y2 and BRAM_Y3 biofilms in presence and absence of 3-Dimensional stress (4% Salinity+ 70°C+ 20% PEG)

The two bacterial strains, BRAM_Y2 and BRAM_Y3, now polyextremophilic bacterial strains were further examined for their plant growth promoting properties. Both the strains grew luxuriantly on Nitrogen deficient medium, thus were nitrogen fixers, solubilizes potassium and phosphate. Both the strains minimally solubilised phosphate BRAM_Y2 at a rate of 9.9 micrograms/ml, and BRAM_Y3 at a rate of 13.5 micrograms/ml in 3 days. (Fig XI(a)). Both the strains also luxuriantly produced the plant hormone IAA both in presence and in absence of a precursor (tryptophan) (Fig XI (b)). The strains also produced Gibberellin, another important plant hormone and ACC Deaminase, the most important enzyme that resists abiotic stress response in plants. The strains were found to produce iron chelating compounds (Siderophores) detected by the orange halo formation by the cell free supernatants when placed in CAS agar medium (Fig XII a and b). The siderophore production by the strains were quantified (Fig XII (c)) and typed, and it was found that BRAM_Y2 produced both the hydroxamate and the catecholate type of siderophores whereas BRAM_Y3 only produced the hydroxamate type. These experiments were followed by the investigation of Biocontrol properties exhibited by BRAM_Y2 and BRAM_Y3. Fig XIII shows the quantitative estimation of the biocontrol cluster of enzymes produced by BRAM_Y2 and BRAM_Y3, which are Catalase, Pectinase, Protease, Beta-1,3-glucanase and peroxidase. Both the bacterial strains were unable to produce HCN or Ammonia as volatile organic compounds. But they excelled in inhibiting a virulent fungus like *Fusarium fujikuroi* (GenBank Accession: OR426451.1) till day 7, in the VOC sealed plate experiment (Fig XIV). Fig XV shows the production of environment sustainability promoting enzymes. Fig XVI is a graphical representation of the two most important vegetative parameters of the plant, “Number of leaves” and the “Plant Height”. The pigment assays also showed significantly better chlorophyll and carotenoid presence in the treated setups in comparison to the untreated control (Fig XVII). Fig XVIII represents the data for two most important reproductive parameters of the plant, “Number of fruits” and “Percent Grain filling” of the fruits. Table II and Table III represent the respective ANOVA tables for the two reproductive parameters, where the null hypothesis was taken to be “Mean of Characteristics are same in both places” and the alternative hypothesis was taken to be “Means being not same in both places”, at a confidence level of 0.05. In both cases the null hypothesis was accepted. Table IV represents the physical parameters of the soil before (initial soil) and after (final soil) cultivation of maize, using BRAM_Y2 and BRAM_Y3 as treatments. Table V represents the Chemical parameters of the soil before (initial soil) and after (final soil) cultivation of maize, using BRAM_Y2 and BRAM_Y3 as treatments.

**Fig XI.**
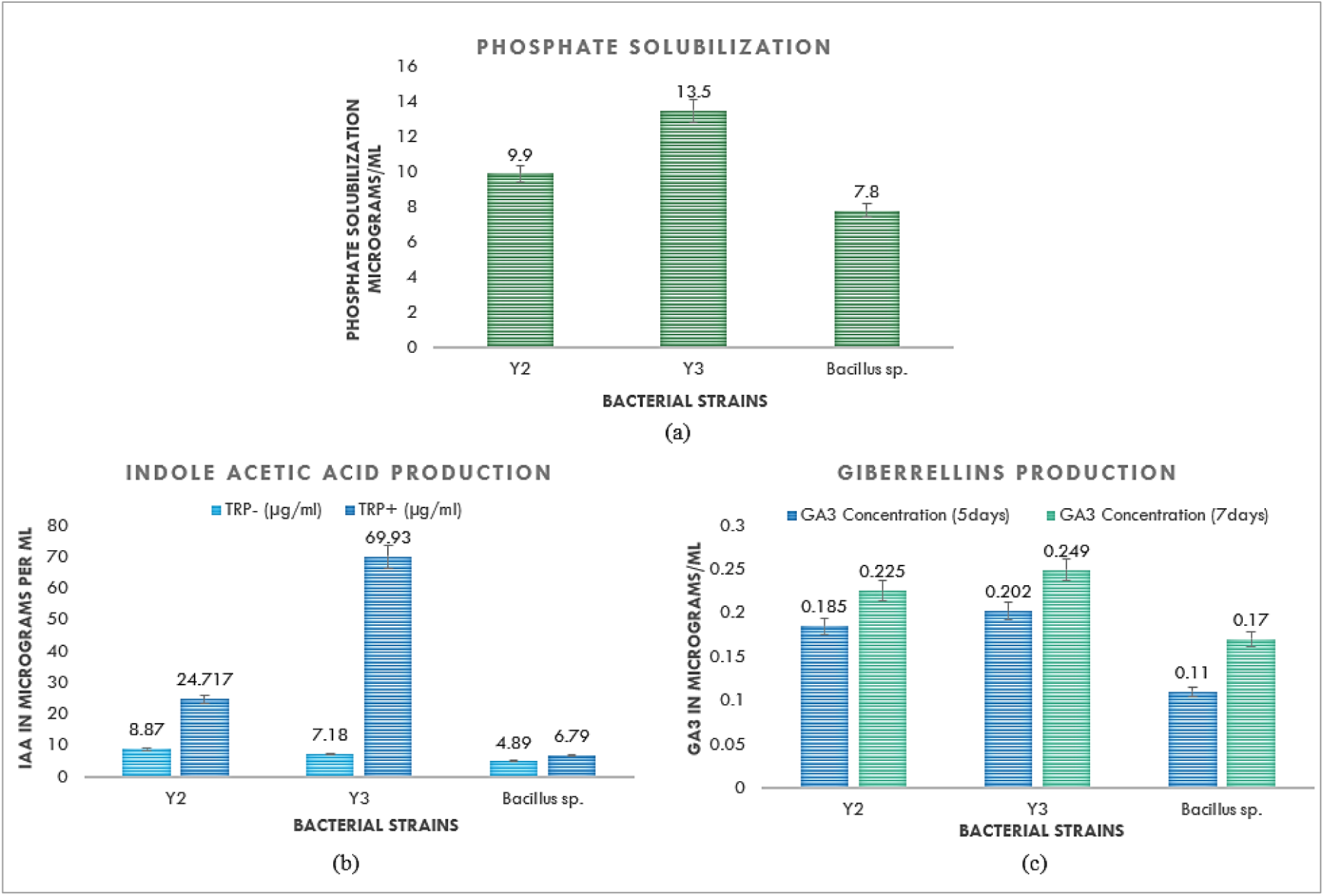
The Plant growth promoting capabilities of BRAM_Y2 and BRAM_Y3 is graphically represented in thisimage. (a) Represents the phosphate solubilising capability of the two strains, (b) represents Indole acetic acid production by the two strains, (c) represents the Gibberellin production by the two strains.

**Fig XII.**
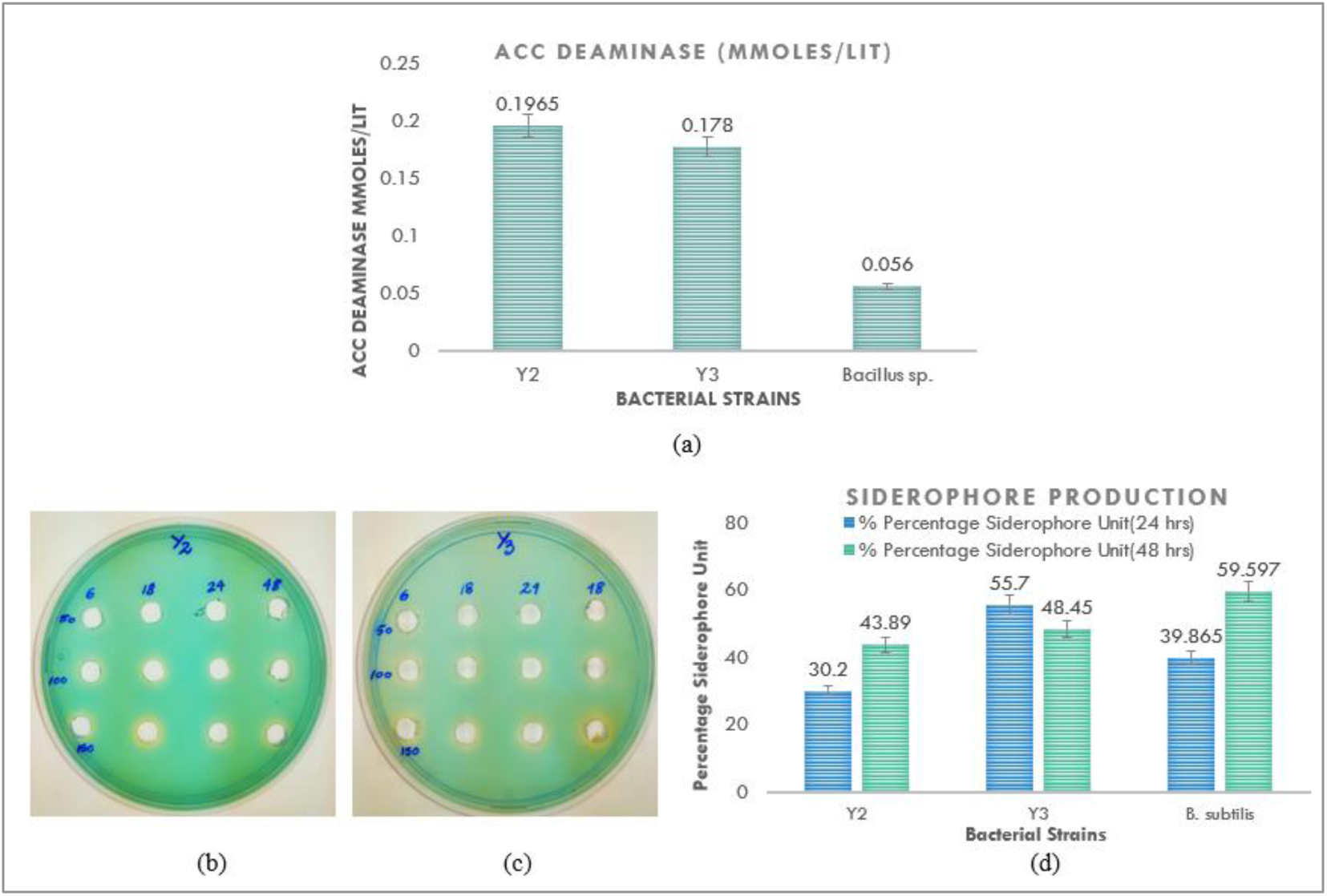
Agriculturally metabolite cluster. (a) (b) and (c) represents the CASAD assay for BRAM_Y2 and BRAM_Y3 respectively where formation of yellow-orange halo indicate presence of siderophores. (d) Represents the Quantified amount of Siderophore production by the two strains BRAM_Y2 and BRAM_Y3.

**Fig XIII.**
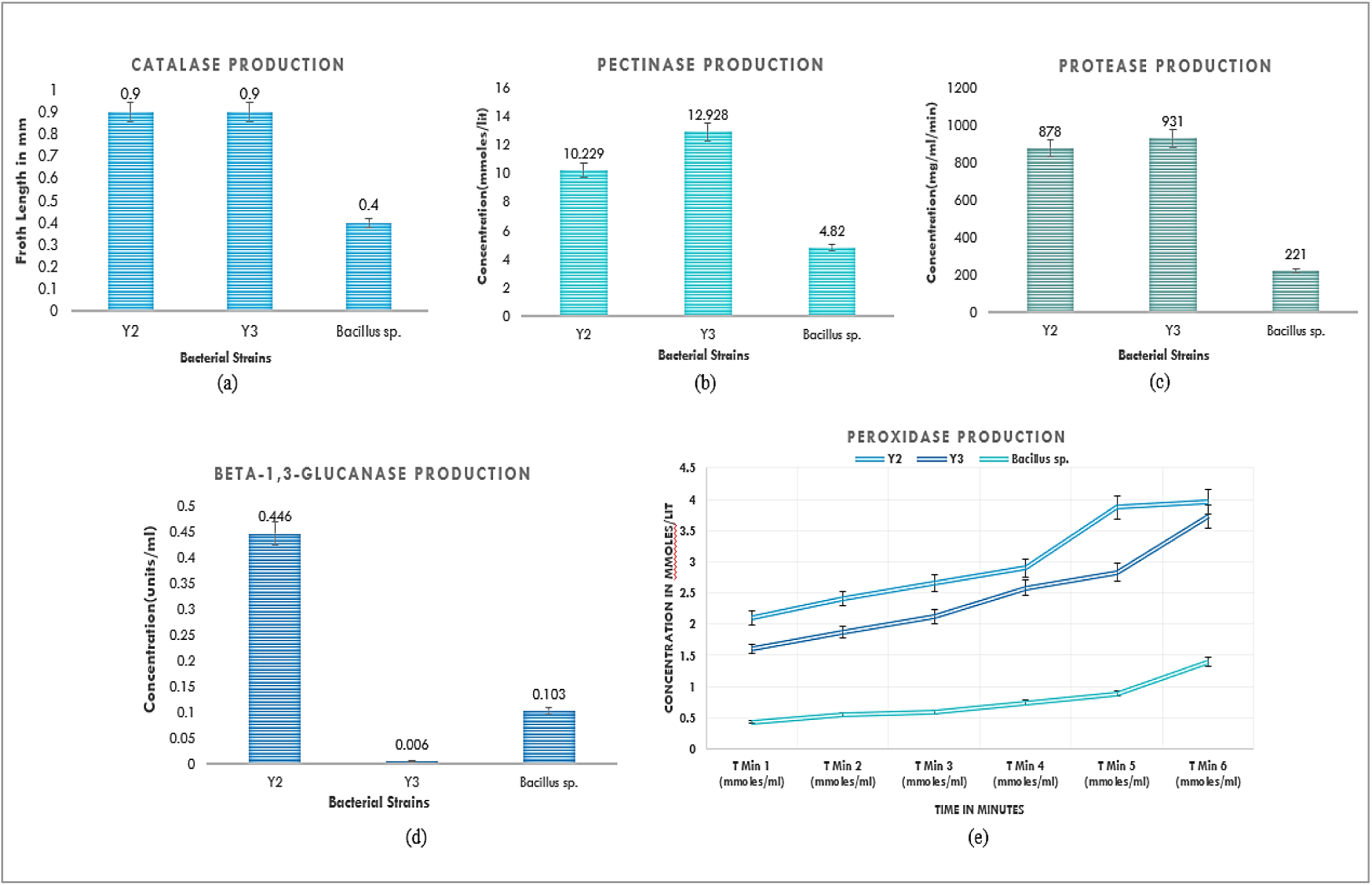
This figure depicts the Biocontrol cluster of enzymes produced by BRAM_Y2 and BRAM_Y3. (a), (b), (c), (d) and (e) Depicts catalase, pectinase, protease, beta-1,3-glucanase and peroxidase production by the two strains respectively.

**Fig XIV.**
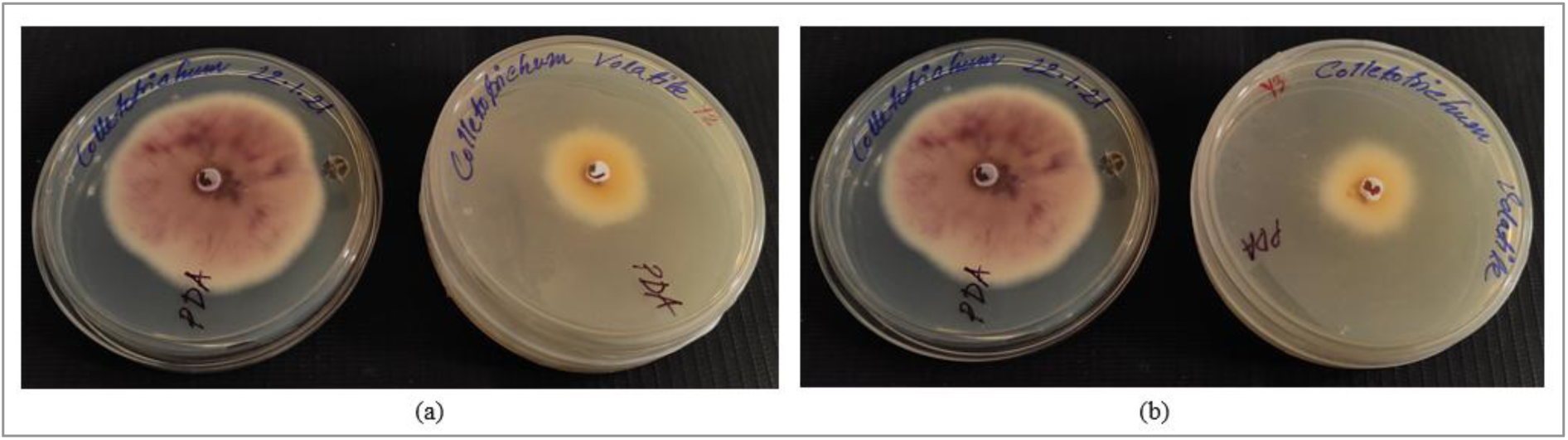
(a) and (b) Sealed Plate for BRAM_Y2 and BRAM_Y3 where it has been tested against *Fusarium fujikuroi (GenBank Accession: OR426451.1*) (The images have been labelled *Colletotrichum sp.* because the preliminary identification of the fungus resembled *Colletotrichum sp* which got rectified after 18srRNA sequencing) by keeping them in same microenvironment with a physical separation to see how it checks the growth of the fungus with the help of its VOCs.

**Fig XV.**
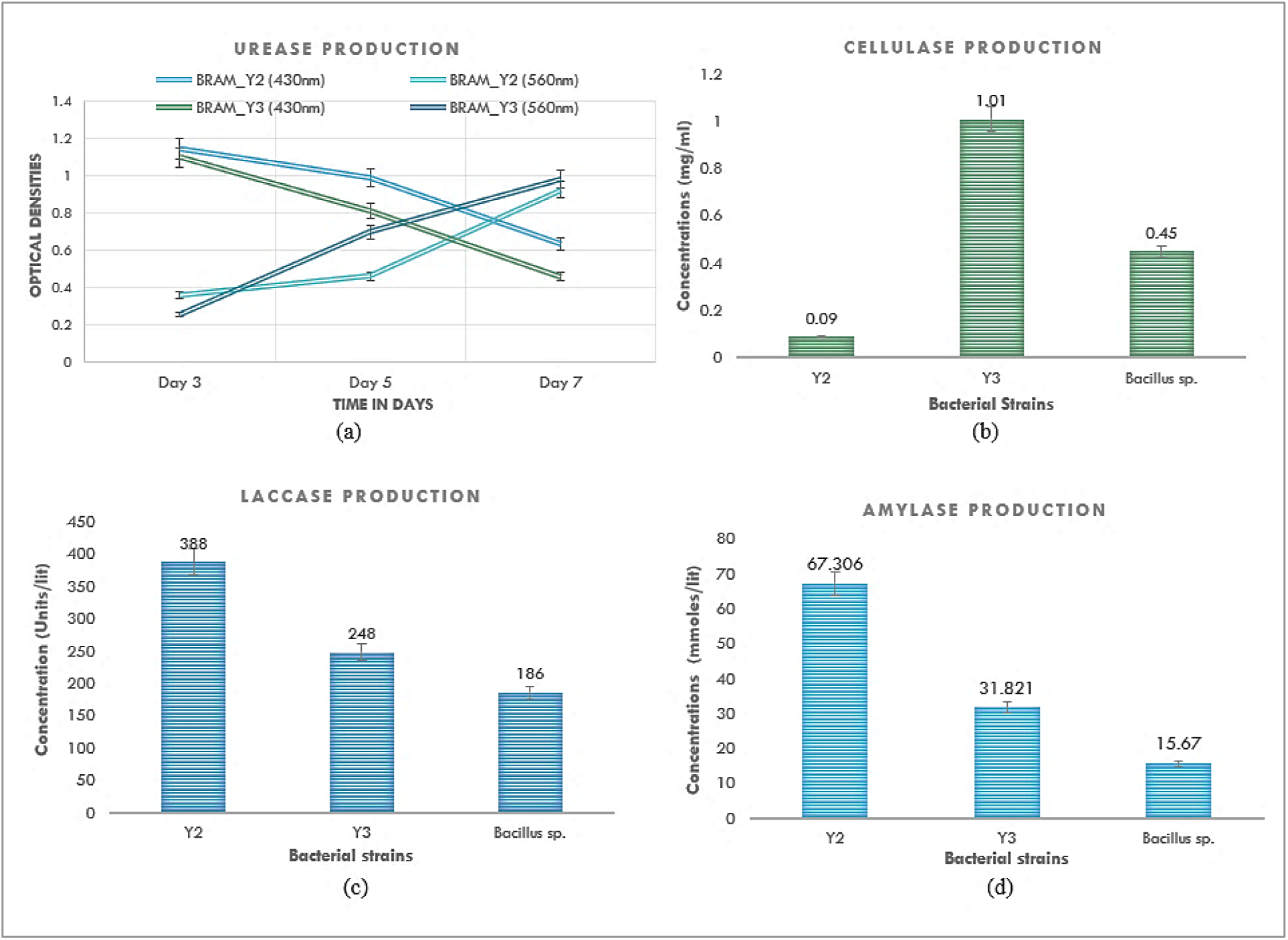
This figure depicts the Environment sustainability cluster of enzymes produced by BRAM_Y2 and BRAM_Y3. (a), (b), (c), and (d) Represents the quantified urease, cellulase, laccase and amylase production by the two strains

**Fig XVI.**
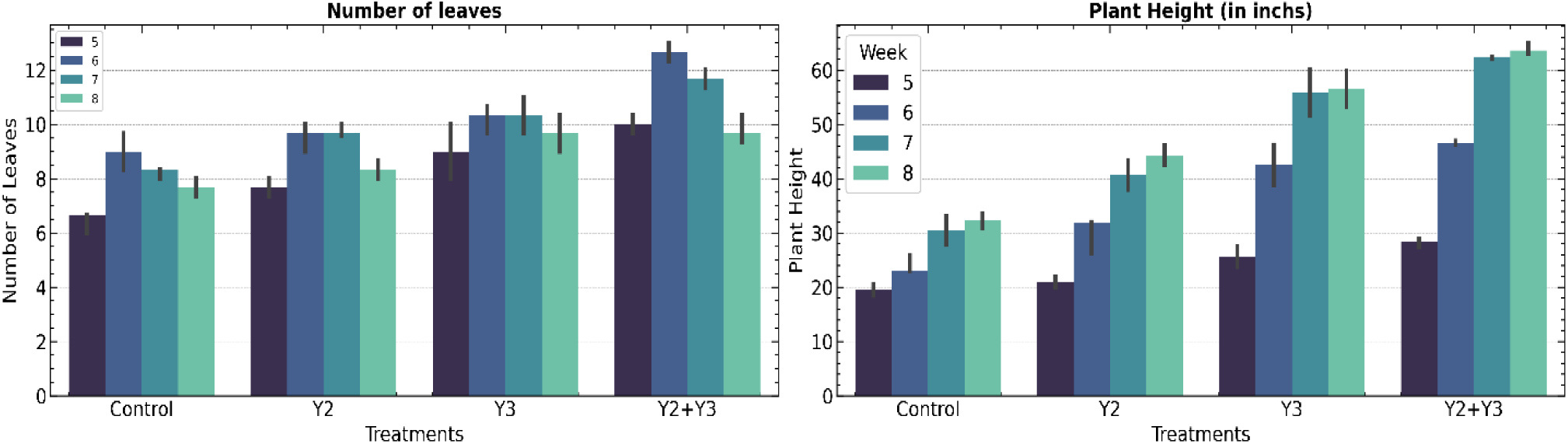
Graphical representation of the vegetative parameters “number of leaves” and “plant height” in the four set-ups, which were control, Y2, Y3 and Y2+Y3

**Fig XVII.**
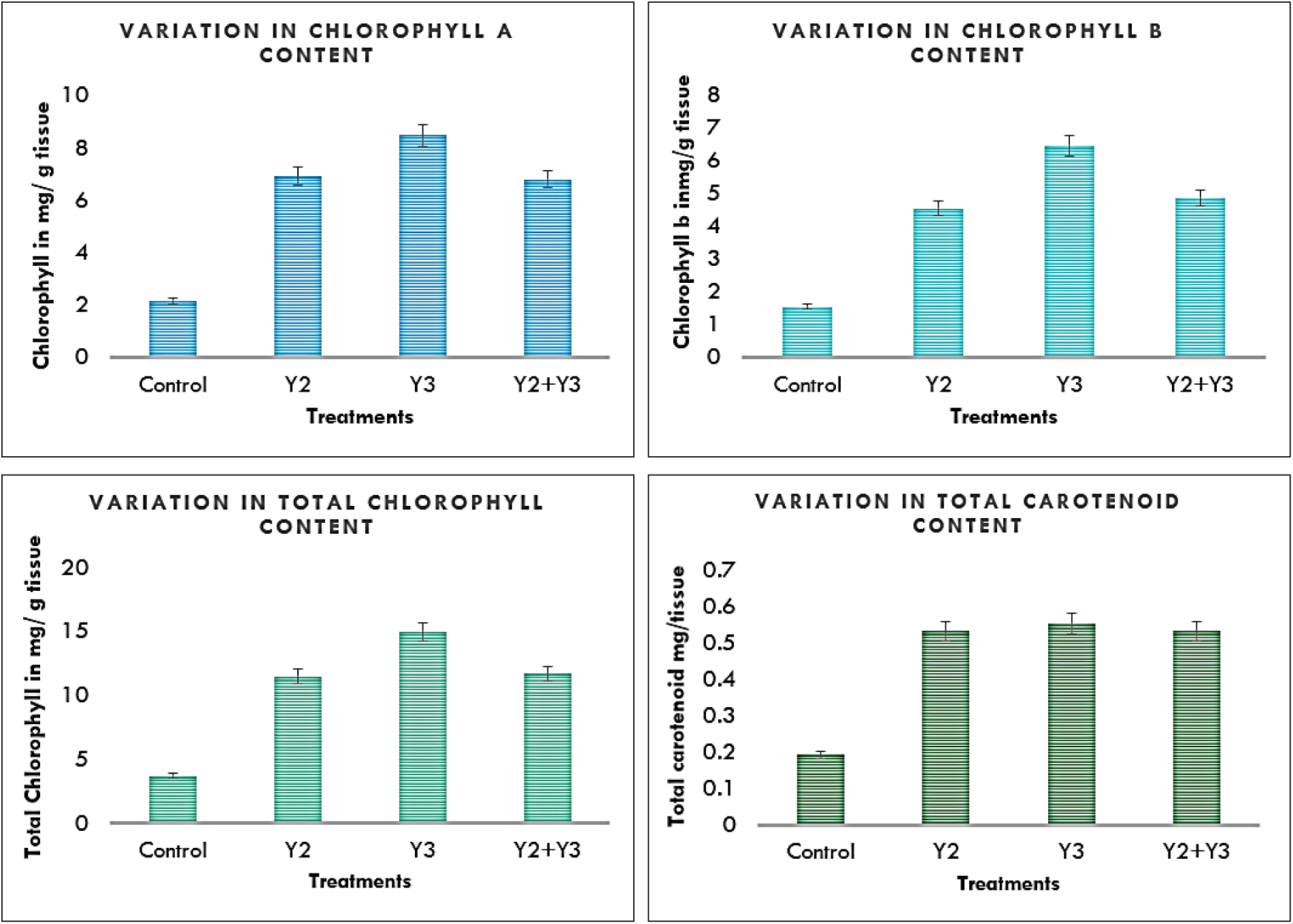
Graphical representation of the variation in the pigments including non-enzymatic antioxidants like carotenoids aft6er treatment application with respect to untreated control.

**Fig XVIII.**
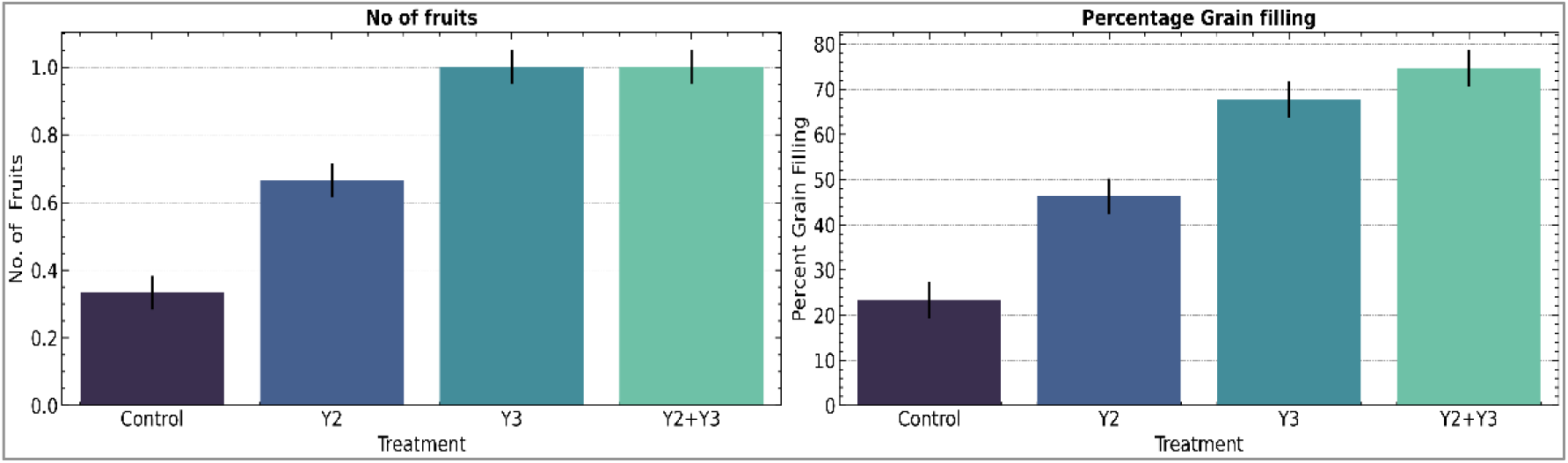
Graphical representation of the reproductive parameters “number of fruits” and “percentage grain filling” in the four set-ups, which were control, Y2, Y3 and Y2+Y3.

**Table II:** ANOVA Table for the “No. of fruits data. ”.

**Independent Variable = Experimental Setup**
|  | Sum of squares | Degrees of freedom | F | PR(>F) |
| --- | --- | --- | --- | --- |
| **Independent variable** | 0.916667 | 3.0 | 1.833333 | 0.219214 |
| **Residual** | 1.333333 | 8.0 | NaN | NaN |

**Decision: Mean of Characteristics are same in both places**

**Table III:** ANOVA Table for the “No. of fruits data”.

| Independent Variable = Experimental Setup |  |  |  |  |
| --- | --- | --- | --- | --- |
|  | Sum of squares | Degrees of freedom | F | PR(>F) |
| Independent variable | 4839.106667 | 3.0 | 1.972698 | 0.196729 |
| Residual | 6541.440000 | 8.0 | NaN | NaN |
| Decision: Mean of Characteristics are same in both places |  |  |  |  |

|  | O.C. | N | P<sub>2</sub>O<sub>5</sub> | K<sub>2</sub>O | Cu | Zn | Fe | Mn | B | Ca | Mg | S |
| --- | --- | --- | --- | --- | --- | --- | --- | --- | --- | --- | --- | --- |
|  | (%) | Kg/ha | Kg/ha | Kg/ha | mg/kg | mg/kg | mg/kg | mg/kg | mg/kg | mg/kg | mg/kg | mg/kg |
| INITIAL SOIL | 0.48 | 439.04 | 110.74 | 215.04 | 5.45 | 8.00 | 15.00 | 10.00 | 0.624 | 881.76 | 607.50 | 6.32 |
| FINAL SOIL | 0.58 | 426.50 | 64.90 | 149.18 | 4.35 | 7.00 | 12.50 | 7.00 | 0.412 | 961.92 | 510.60 | 3.20 |

**Table IV:** Physical Parameters of the Soil before and after application of the Bacterial strains BRAM_Y2 and BRAM_Y3 as treatment during field trail on *Zea mays* L.

|  | pH | E.C. | CEC | Sand | Silt | Clay | Texture | BD | WHC | Pore | Moisture | Humic |
| --- | --- | --- | --- | --- | --- | --- | --- | --- | --- | --- | --- | --- |
|  |  | dsm<sup>-1</sup> | meq/100gm | (%) | (%) | (%) | of Soil | gm/cc | (%) | Space (%) | (%) | Acid (%) |
| INITIAL SOIL | 7.88 | 0.16 | 14.80 | 44.40 | 36.00 | 19.60 | Loam | 1.39 | 34.93 | 43.26 | 7.60 | 6.00 |
| FINAL SOIL | 8.26 | 0.11 | 11.76 | 46.40 | 36.00 | 17.60 | Loam | 1.32 | 36.44 | 44.59 | 17.40 | 4.00 |

**Table V:** Chemical Parameters of the Soil before and after application of the Bacterial strains BRAM_Y2 and BRAM_Y3 as treatment during field trail on *Zea mays* L.

## Discussion and Conclusion

The bacteria *Mesobacillus subterraneus* was first reported by Kanso. S et al, 2002, isolated from an Australian thermal aquifer. *Brevibacillus parabrevis* was first studied by Takagi et al in the year 1993 and then Hooda, R, 2018. Both the bacterial species are rare in nature and not much studies correspond to their characterization and application. In this investigation, *Mesobacillus subterraneus* BRAM_Y2 and *Brevibacillus parabrevis* BRAM_Y3 colony characteristics were vividly studied, the characteristic pigmented yellowish orange colour of the BRAM_Y2 colonies and the opaque white colonies of BRAM_Y3 with the characteristic pearl-like shine was reported for the very first time (Fig I). Unlike most *Bacillus* sp. both the bacterial strains didn’t form endospore.

Both the bacterial strains also exhibited numerous extremophilic properties and produced considerably high amount of biofilm as compared to the control strain (Fig III). The bacterial strains were found to give exuberant growth in temperatures ranging from 20°C to 70°C (Fig IV). Though both of them survived in humongous temperature range of −20°C to 120°C make them both qualified to be called thermophiles and psychrophiles (Blochl. E et al., 1995). This was followed by their tolerance of salinity. The two strains were found to tolerate 4% of salt (Fig V(a)), which characterized them as moderate Halophiles (Barrozzi. A et al., 2018). The results for pH tolerance exhibited a huge range of pH tolerance of 1-12 in both BRAM_Y2 and BRAM_Y3 though the pH optima for BRAM_Y2 was seen to be at pH 12 and for BRAM_Y3 at pH 10. The results indicate the existence of both Acidophilic and alkalophilic nature of the two bacterial strains (Fig V (b)). The control strain of lab *Bacillus* survived through the pH of 5-9, with the optima at pH 7(Schleper C et al., 1995). The two test strains of BRAM_Y2 and BRAM_Y3 tolerated water activities(a_w_) of 0.85-0.416, decreased by the presence of Polyethylene glycol 6000 in the medium (Fig V c). This categorised the two strains as Xerophiles (Lebre et al., 2017). The two strains under investigation were then subjected to Ultraviolet light, of 254nm wavelength and 11 microwatt/cm^2^ for 6 hours, which made the total dosage of UV of them to be 3.9J/m^2^, 15 mis of which is expected to kill any bacterial strain. But astonishingly, the bacterial strains gave faster growth compared to their normal growth in the presence of UV (Fig V d). This property classified them as radio-resistant microbes (Pullerits et al., 2020;). The tolerance of the two bacterial strains BRAM_Y2 and BRAM_Y3 to 200ppm of Iron, 300ppm of Arsenic, 20ppm of silver and 10ppm and 15ppm of mercury respectively designated them to be metallophiles as well (Fig VI) (Golyshina OVet.al, 2000). Keeping in mind the excellent biofilm forming capability of the bacteria, the increase in biofilm formation with respect to single dimension was measured (Fig VII b).

To conclusively demonstrate Polyextremophily in BRAM_Y2 and BRAM_Y3, they were subjected to 16 combinations of two-dimensional stress and 9 combinations of three-dimensional stress, where not only they grew luxuriantly in most of the combinations, they also showed extremely high biofilm formation, the significance of which is discussed in the latter half of this discussion (Fig VIII and IX) (Roy et al., 2022).

The two bacterial strains BRAM_Y2 and BRAM_Y3 now better discernible as polyextremophilic strains, were explored for their plant growth promoting abilities. Both the strains solubilize phosphate at minimal levels (Fig X a). Plant growth regulator, Indole acetic acid and gibberellins which are immensely important for plant growth and development are produced by both the strains in significant amounts (Gusmiaty et al., 2019). IAA is produced by BRAM_Y2 even in absence of its precursor, tryptophan, and in very minimal amounts by BRAM_Y3 (Fig X b and X c).

The fourier transform infrared spectral scans in the images represent the different molecular bonds corresponding to classes of compounds such as lipids, proteins, amino acids, molecular oxygen bonds and so on. The decrease in transmittance corresponds to increase in absorbance and therefore increase in the concentrations of certain classes of compounds. The fourier transform infrared spectral scans of the extracted biofilms of the 2 bacterial strains in presence and absence of a three-dimensional stress exhibit distinct differences with respect to peak hieghts (concentrations) and peak shifts. The three-dimensional stress combination used in this experiment and the rest of the experiment here onwards was selected on the basis of the overall maximum growth observed by all 2 bacterial strains in the aforesaid combination of Temperature 70 °C + 4% NaCl+ 20% PEG. Three categories of peaks were mainly observed in the FTIR scans of the biofilm extracts, 3000-3500cm^-1^that corresponds to lipids and proteins, 1500-1700 cm^-1^that corresponds to amide bonds (I and II) and C=C conjugates and lastly 500-600 cm^-1^ which can be molecular oxygen or carbon and halogen bonds (tetrahedral). The main observable change that occurred in 3D-stress and no stress conditions was that there was a huge increase in the peak heights and therefore the corresponding concentrations of the various compounds. The other change that could be observed was the slight shifts in the peaks though they were present in range of the same class of compounds which might correspond to change in forms of the compounds in stressed conditions. These observations clearly indicate a crucial role played by bacterial biofilms in protecting them in times of abiotic stress conditions.

ACC Deaminase is a very important enzyme produce by bacteria in the plant rhizosphere to combat abiotic stress response in plants. ACC is the precursor of plant stress hormone Ethylene, secreted in the plant rhizosphere by the roots in times of any abiotic stress such as salinity, drought, heavy metals etc. Bacterial ACC Deaminase dissociates ACC into ammonia and α-keto butyrate. Thus, inhibiting the formation of ethylene, which when formed, induces heavy damage to the plant physiology (Glick, 2013; CarmenOrozco-Mosqueda et al., 2020). the two bacterial strains produce notable amounts of ACC Deaminase to support plant growth in abiotic stress conditions (Fig XII a).

There can be two types of Siderophores, hydroxamate and catecholate, that help in iron acquisition are also produced by the bacterial strains, that can rescue the plants growing in low iron conditions. They might also have roles in biocontrol of phytopathogens, by limiting iron for them in the environment (Ahmed et al., 2022). Both the bacterial strains produced considerable amounts of siderophore in both 24hrs and 48hrs (Fig XIId). Enzymes, required for Biocontrol of plant pathogens such as catalase, pectinase, protease, β-1,3-glucanase and peroxidase are produced by the two strains, Though, BRAM_Y3 is incapable of producing β-1,3-glucanase. These enzymes for a part of the plant’s Induced Systemic Resistance pathway (Fig XIII) (Heil and Bostock, 2002). In addition to liquid exudates, the bacterial strains BRAM_Y2 and BRAM_Y3 are also capable of producing certain unknown volatile organic compounds (unknown, as neither of them produced ammonia or hydrogen cyanide gasses on experimentation) that efficiently inhibits the growth of a well-known plant pathogen, *Fusarium fujikuroi* (GenBank Accession: OR426451.1) (Fig XIV) (Ruangwong et al., 2021). All the aforementioned experiments validated the fact that BRAM_Y2 and BRAM_Y3 have great potential as plant growth promoting agents.

Further exploration of the properties of the two bacterial strains were done on their ability of producing enzymes, related to environmental sustainability. Urease, an enzyme that degrades urea to release carbon dioxide and ammonia is one of such important enzymes. Excessive use of Urea in the agricultural fields culminates in a huge residual toxicity as the plants can only utilise a very small percentage of the supplied urea. The bacterial urease removes this excess urea from the soil, in turn reducing its toxicity (Inamdar et al., 2022). Both the bacterial strains BRAM_Y2 and BRAM_Y3 produce impressive amounts of urease which would contribute a lot to soil health (Fig XV a). Ligno-cellulytic enzymes such as laccase, amylase, cellulase etc. play an essential role to recycle the lignocellulosic wastes in the agricultural fields and increase the organic carbon content of the soil, therefore improve the soil quality. The test bacterial strains also produce satisfactory amounts of these enzymes again contributing to soil health (Fig XV b, XV c and XV d).

With all these properties at hand, these bacteria were applied on test crop *Zea mays* L to check for their in-vivo potential as PGP. The two bacterial strains BRAM_Y2 and BRAM_Y3 individually and together in a consortium showed impressive results in terms of both reproductive and vegetative with respect to the control. The consortium performed best in the context on vegetative parameters (Fig XVI). In terms of number of fruits BRAM_Y3 and the consortium of BRAM_Y2+ BRAM_Y3 performed the best, though BRAM_Y2 was better that control it was not as good as the rest. The pigments like Chlorophyl and carotenoids, which is also a non-enzymatic antioxidant, the treatments showed remarkably higher values than that of the untreated control (Fig XVII). The increased carotenoids can help the plant protect itself from different biotic and abiotic stresses as well (Swapnil et al., 2021). In terms of Grain filling, the consortium of the two bacterial strains showed the best results (Fig XVIII). Soil analysis before and after cultivation of *Zea mays* L with bacterial treatments showed an increase in soil organic carbon, pore space and moisture. On the other hand, there was a very slight decrease in soil Nitrogen, because though one of the most utilised nutrients, due to the presence of Nitrogen fixers as treatment, the change was very small. A significant decrease in the levels of potassium and phosphate due to solubilization by bacteria and utilization by plants. Similarly decrease in the levels of Zinc, Iron, Manganese, Magnesium and sulphur was also observed (Table IV and Table V).

According to Maitra et al., 2022, biofilm plays an integral role in the plant growth promoting properties of a bacteria. Higher the biofilm formation, better is the root colonization of the rhizospheric bacteria, and better are the production of plant growth promoting metabolites. Thus, as stress increases the biofilm, the plant growth promoting properties can remain unhampered or get enhanced for that matter. The experimental evidences found in this study and in Roy et al., 2022 that stress application has a significant impact on increased biofilm formation. Now if there is a direct relation between biofilm formation and plant growth promotion activity which was again substantiated by Maitra et al., 2022, then the two bacterial strains *Mesobacillus subterraneus* BRAM_Y2 and *Brevibacillus parabrevis* BRAM_Y3 as Plant Growth Promoters in harsh and depleted soils can open up new horizons for organic cultivation in extreme environments like saline soils, alkaline or acidic soils, arid and semi-arid regions and even in areas where the temperatures have risen significantly or in the past few decades due to the global phenomena of climate change.

## Ethical Approval

Not applicable.

## Consent to Participate

Not applicable.

## Consent to Publish

Not applicable.

## Author’s Contributions

BR- conceptualization, Research, preparation of the original draft, DM- conceptualization, research, review of the original draft, AC- design of machine learning-based models and data analysis, PB: conceptualization & supervision IC- conceptualization, & supervision, AKM- planning, conceptualization, supervision and acquisition of fund, JG- supervision and guidance

## Acknowledgement

We would like to thank Vivekananda Institute of Technology, Nimpith for facilitating the tests on soil parameters. We would like to thank Eurofins, Bangalore, for facilitating the 16S rRNA studies. We would like to thank Dr. Satadal Das, B.K. Roy Research Centre, Peerless Hospital Kolkata for his inspiration for the work.

## Availability of Data and Materials

All materials are reported in the text, all the data collected are reported in the manuscript

